# Evolutionary *PTEN* gene divergence underpins the remodeling of plant vacuolar compartments

**DOI:** 10.1101/2022.01.18.476728

**Authors:** Bojan Gujas, Chloe Champeyroux, Anna Hunkeler, Emilija Robinson, Noel Blanco-Touriñán, Tiago Miguel Dias Cruz, Matthias Van Durme, Moritz K. Nowack, Antia Rodriguez-Villalon

## Abstract

Membrane fusion and fission are fundamental processes in sustaining cellular compartmentalization. Fission of a lipid bilayer requires a furrow formation that brings membranes in close proximity prior to a contiguous membrane cleavage. Although plant ancestors abandoned cleavage furrow-mediated cytokinesis more than 500 million years ago, here we show that plants still employ this mechanical principle to divide embryonic vacuoles. The evolutionary divergence in PHOSPHATASE AND TENSIN HOMOLOG DELETED ON CHROMOSOME TEN (PTEN) enzymes was required to coordinate this process, as Arabidopsis loss-of-function *pten2a pten2b* mutants contain hyper compartmentalized embryonic vacuoles. In contrast, *PTEN2* overexpression hinders lytic and secretion cellular pathways downstream of TGN in xylem cells. These processes are critical for the formation of secondary cell walls in xylem cells and depend on a poorly characterized and evolutionarily novel N-terminal domain in PTEN2s. The *PTEN2* subfamily appeared with the emergence of the *Phragmoplastophyta* clade, when vacuolar compartments enlarged and cleavage furrow-mediated cytokinesis became extinct. Together, our work suggests that the evolutionary innovation of the PTEN family is conserved across terrestrial plants and central to vacuolar remodelling.

## INTRODUCTION

Cell compartmentalisation is a basic organisational principle of eukaryotic life that separates variety of partitions within the cell to generate multiple different metabolic environments. Communication between endomembrane compartments occurs by a tightly regulated interplay of membrane fission and fusion^1, 2^. Although the molecular players involved in membrane fission can vary across living kingdoms, a common mechanistic requirement is to bring two membranes into close proximity^1, 3^. Generally, this energetically costly membrane bending is followed by a furrow formation at the cleavage site that progresses centripetally until a neck structure is generated on which cleavage proteins can act^3, 4^. The membrane bending requires dynamic changes in its physiochemical properties. For example, the activation of phosphatidylinositol 3-kinase (PtdIns3-kinase) to produce phosphatidylinositol 3,4,5-trisphosphate (PtdIns[3,4,5]P3) was shown to be a prerequisite for membrane ruffling in mammalian cells as well as for the leading membrane bending during neutrophil cells or amoeba chemotaxis^5, 6^. Membrane relaxation is achieved by PtdIns[3,4,5]P3 dephosphorylation at the 3’ position by PHOSPHATASE AND TENSIN HOMOLOG DELETED ON CHROMOSOME TEN (PTEN) activity from the rear sides of the motile membrane^7^. Interestingly, the phosphoinositides species phosphorylated at 3’ positions are scarce in plants, while PtdIns[3,4,5]P3 has never been detected^8, 9^. Yet, the *Arabidopsis thaliana* (Arabidopsis) genome encodes three *PTEN* homologs split in two subfamilies: *PTEN1*, and *PTEN2* comprised of the paralogs *PTEN2a* and *PTEN2b* (*PTEN2s*)^10^. The need for expansion of *PTEN* genes in plants is unclear to date.

In the green lineage, cytokinesis mechanism evolved from ancestral centripetal cleavage, still occurring in algal *Streptophyta* clade, to centrifugal fusion model where vesicles fuse to the growing septum membrane called cell plate^11, 12^. Membrane donors here are guided by the evolutionary novel structure – phragmoplast, thus all organisms using this model of cytokinesis are collectively termed *Phragmoplastophyta*, including algal division Charophyta as well as all land plants^12^. In addition to discontinuation of a cleavage furrow for cytokinesis, plant cells enlarged their vacuolar compartment in comparison to unicellular green algae in the protist clade^13^. These large compartments became essential for land plant cells’ viability as they provide plant hydrostatic skeleton in a form of turgor pressure^14^. Additionally, vacuoles display a vast array of cellular functions^13^. Vegetative tissues contain lytic vacuoles, which are acidic compartments abundant in hydrolases critical for ion homeostasis and lytic degradation^13^. In contrast, protein storage vacuoles (PSVs) accumulate protein reserves that fuel plant development during germination and are characterized by a neutral pH^13^. Recently, *PTEN2a* was implicated in vacuolar trafficking in Arabidopsis^15^ - consistent with the reported PTEN2s substrate preference to be PtdIns3P, typically residing in membranes of lytic compartments^9, 10, 15^. The latter poses the question whether plant *PTEN2s* may have a role in remodelling vacuolar membrane (tonoplast) instead of the plasma membrane as reported in other eukaryotes. However, how voluminous vacuolar compartments divide in plants is still a matter of debate^16^. Recent 3D models of vacuoles suggested that the compartments that appear fragmented in 2D images are actually not physically separated but rather form a tubular network^17–19^. Thus, many previously reported examples of a vacuole fragmentation have to be revalidated in order to gain better understanding of vacuolar membrane dynamics.

Here, we followed the stepwise conversion of large embryonic vacuoles (EVs) into smaller PSVs in Arabidopsis. Our 3D reconstructions of EV remodelling revealed vacuolar division by a process morphologically resembling the progressing cleavage furrow division. Moreover, we show that PTEN2 enzymes are essential to coordinate membrane tubularization at EV division initiation sites and cleavage furrow progression. In *pten2a pten2b* double mutants EVs become hyper-compartmentalized instead of fragmented. On the contrary, overexpression of *PTEN2s* prevents the fusion of trans-Golgi network (TGN)-derived small vacuoles to the central vacuole that does not enlarge but stays tubular. This phenomenon was cell type specific and predominantly occurring in xylem tissues. Aberrant cell trafficking affected both vacuolar and secretory pathways, essential for xylem tissue maturation. Notably, *PTEN2* function in remodelling vacuolar architecture depends on their poorly characterized N-terminal domain, that evolutionarily appeared in the *Phragmoplastophyta* clade, coinciding with vacuolar enlargement and loss of cleavage furrow-mediated cytokinesis^20^. Thus, it seems plausible that *PTEN2s* evolved to provide molecular support to preserve this ancient model of membrane fission to modulate the biggest plant cell compartment - the vacuole.

## RESULTS

### Embryonic vacuole division involves cleavage furrow formation and requires PTEN2 activity

During plant embryogenesis, EV become transformed into numerous small PSVs prior seed desiccation^21, 22^. To assess whether this process entails vacuolar fragmentation, we followed the dynamics of previously described vacuole marker *TONOPLAST INTRINSIC PROTEIN (TIP) 3;2* during Arabidopsis embryo development. Expressed from a native *TIP3;2* promoter fragment, this marker is detectable from the late heart or early torpedo stage of Arabidopsis embryogenesis onwards. TIP3;2-GFP first accumulates in the endoplasmic reticulum (ER) around the nucleus and near the plasma membrane, similar to the V-PPase VHP1/AVP1 tonoplast marker at the same stage (stage I in Fig. 1a Extended Data Fig. 1a). In succeeding stages of embryo development, the first pre-EV can be observed in addition to the ER signal (stage II). Next, the TIP3;2 accumulation becomes restricted to the tonoplast of pre-EVs that undergo fusion, evident by the hollowed spaces bridging two vacuoles at the fusion site (stage III in Fig. 1a and Extended Data Fig. 1b). As a result of these homotypic fusions, larger EVs are generated (stage IV in Fig. 1a), followed by the onset of the typical PSV autofluorescence in the successive stages. Next, the large EV starts dividing, evidenced by the symmetrical tonoplast invaginations towards the vacuolar lumen (stage V in Fig. 1a). By performing 3D reconstructions based on maximal projections and surface rendering we could observe formation of a cleavage furrow-like structure in the region where the tonoplast is contracting around the incipient separation site (Fig. 1b). As one EV splits in multiple PSVs, various cleavage progression stages can be observed in a single EV. Stage VI is characterized by the final fragmentation of EV into multiple PSVs (Fig. 1a). During the process of EV division in some samples we noticed the presence of small vacuoles that may be the membrane source necessary for tonoplast invagination during furrowing or suggest an additional alternative pathway of vacuole fragmentation (Extended Data Fig. 1c).

**Figure 1:**
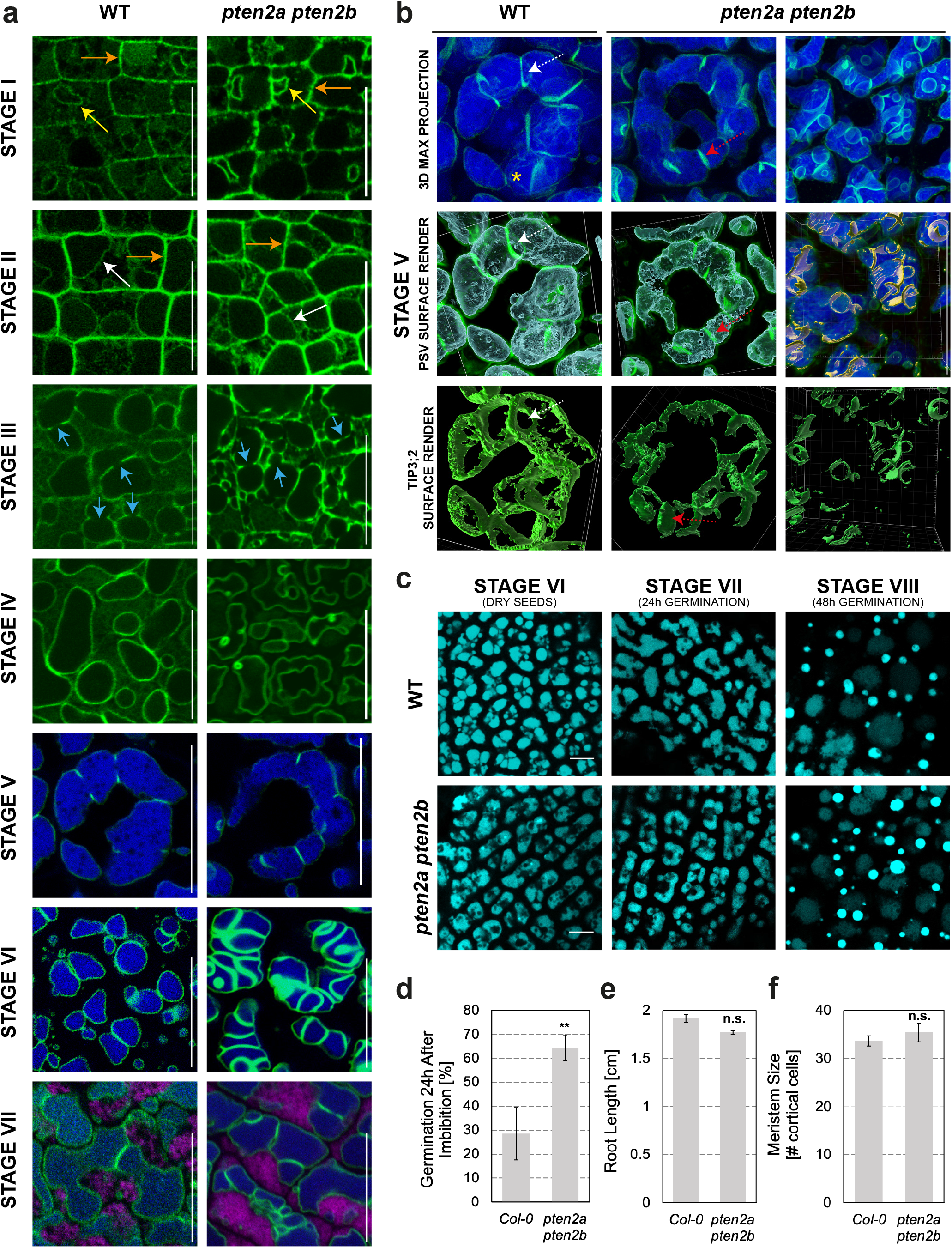
Embryonic vacuole division involves cleavage furrow formation and requires PTEN2 activity. **a-h**, Representative confocal images of vacuoles in epidermal cells in Arabidopsis embryos extracted from green siliques (stages I-III), yellow siliques (stages IV-V), dry seeds (VI), germinated for 24h (stage VII) or 48h (stage VIII). Tonoplast (vacuolar membrane) decorated by *TIP3;2-GFP* (in green). **a**, Comparative seven stages of TIP3;2 dynamics between wild type and *pten2a pten2b*. At the onset of its expression, TIP3;2 accumulates in endoplasmic reticulum close to plasma membrane (orange arrows), or follow nuclear shape (yellow arrows). First tonoplast is visible in stage II marked with a white arrow. Small pre-embryonic vacuoles fuse in stage III (fusion sites are marked with blue arrows). Please note that image of EVs in *pten2a pten2b* in stage IV is slightly advanced than in wild type. In stages V-VII PSV exhibit autofluorescence in blue part of the spectrum (represented blue in images). Magenta shows autofluorescence in red part of the spectrum. **b**, EV division starts by tonoplast invagination at division site visible as a circle (white dashed arrows). In addition to the lack of synchronicity in tonoplast invaginations, in *pten2a pten2b* double mutants, membrane unilaterally ingress (red dashed arrow) and can reach the other side of the vacuole, however the division does not occur. Possible cause is the absence of membrane bending into hourglass shape as visible in WT (yellow asterisk). The ingrown membrane possibly rolls into a cylindrical shape as visible in 3D reconstructions of the late-stage V in double mutants. **c**, Representative confocal images of storage vacuoles’ autofluorescence in epidermal cells of Arabidopsis embryos extracted from dry seeds and germinating seedlings in wild type and *pten2a pten2b* mutants. Scale bars represent 20μm. **d**, Germination 24h after imbibition in indicated genotypes. Error bars represent SE. n>500. **e-f**, Quantification of root length (**e**) and meristem size (**f**) in 6 days old seedlings of the indicated genotypes. n>30 (e) or n=10 (f). All error bars represent standard error. n.s., not significant; ∗∗p < 0.01.

To determine the molecular mechanisms underpinning embryonic vacuolar division in plants, we decided first to explore whether PtdIns3P metabolic enzymes are involved in the regulation of this process. In budding yeast, a local enrichment of PtdIns3P at the neck occurs before membrane fission and it is required to stabilize membrane invaginations^23^. Detailed examination of *pten2a pten2b* mutants revealed a crucial difference in the progression of EV division in stage V in comparison to wild type embryos (Fig. 1a). In *pten2a pten2b* embryos the tonoplast inward invaginations are not coordinated, and often asymmetric unilateral. The invaginated membrane can be seen in 3D as individual sheets growing inwards; and even when these sheets meet the other side of the vacuole, a ring-shaped cleavage furrow-like structure will be absent or incomplete (Fig. 1b). The membrane sheet would in later stages roll in on itself and form a cylindrical shape, or sometimes a sphere that may eventually pinch off. At the end, instead of fragmented, EVs of *pten2a pten2b* mutants appear hyper- compartmentalized with their vacuolar lumen crisscrossed with several membranes (stage VI in Fig. 1a). During seedling germination, the homotypic fusion of PSVs (stage VII in Fig. 1a and 1c) generates the central lytic vacuolar compartment, a process that can be scored by the gradual disappearance of the typical PSV autofluorescence by 48h after seed imbibition. Surprisingly, lytic vacuole formation appeared unaffected in *pten2a pten2b* double mutants, although mutant seedlings germinated slightly faster than wild type (Fig. 1d and Extended Data Fig. 1d). Yet, this process does not translate in a faster post-embryonic growth, as manifested by a similar root growth and underlying meristematic activity (Fig. 1e-f).

### PTEN2s overexpression prevents xylem cell differentiation

To further elucidate the potential role of PTEN2s in vacuolar remodeling and its effects on plant development, we decided to analyze the impact of altered *PTEN2* levels. As, the constitutive *PTEN2*b overexpression from UBQ10 promoter was lethal, we employed the estradiol inducible system to assess its short-term overexpression effects in relevant cell types (Extended Data Fig .1e). During germination, the consumption of PSV reserves was proposed to occur primarily in vascular cells, most likely to allow for the development of a functional vascular system before the amino acids are mobilized from other parts of the plant body^24, 25^. Hence, we focused on xylem cells. To become conductive units, xylem cells undergo a complex developmental process that encompasses the reinforcement of the cell wall (SCW) and a vacuolar- driven programmed cell death (PCD)^26, 27^. The formation of fully differentiated xylem cells with lignin-reinforced SCWs can already be observed 48h after seed imbibition (Extended Data Fig. 1f). While *pten2a pten2b* double mutants showed neither acceleration nor delay in xylem vessel differentiation, seedlings overexpressing *PTEN2b* showed significant defects in xylem maturation (Extended Data Fig. 1f). Interestingly, the xylem differentiation inhibitory effect had only overexpression of *PTEN2* paralogs but not *PTEN1* isoform (Fig 2a-b), suggesting a functional evolutionary divergence between orthologs. In the further text we will mainly focus on *PTEN2b* overexpressing lines (*PTEN2b_ox_*), as the results obtained in either *PTEN2a* or *PTEN2b* overexpressing lines were very redundant. The phenotypical effects of *PTEN2_ox_* were dosage dependent and sensitive to the duration of induction (Extended Data Fig. 1e and 1g-h). The induction with a lower estradiol concentration (0.2μM) for shorter time (24-48h) did not affect much the overall root growth, but inhibited xylem differentiation completely or allowed only individual cells to differentiate (xylem islands) (Fig 2b). Higher expression detected in independent transgenic lines or achieved by estradiol induction in higher concentrations (2μM) significantly shortened the root length but reverted xylem phenotypes almost to normal (Extended Data Fig. 1e and 1g-h). The latter was scored mainly in distal root parts where the tissue was longer exposed to *PTEN2b* induction agent prior differentiation.

**Figure 2:**
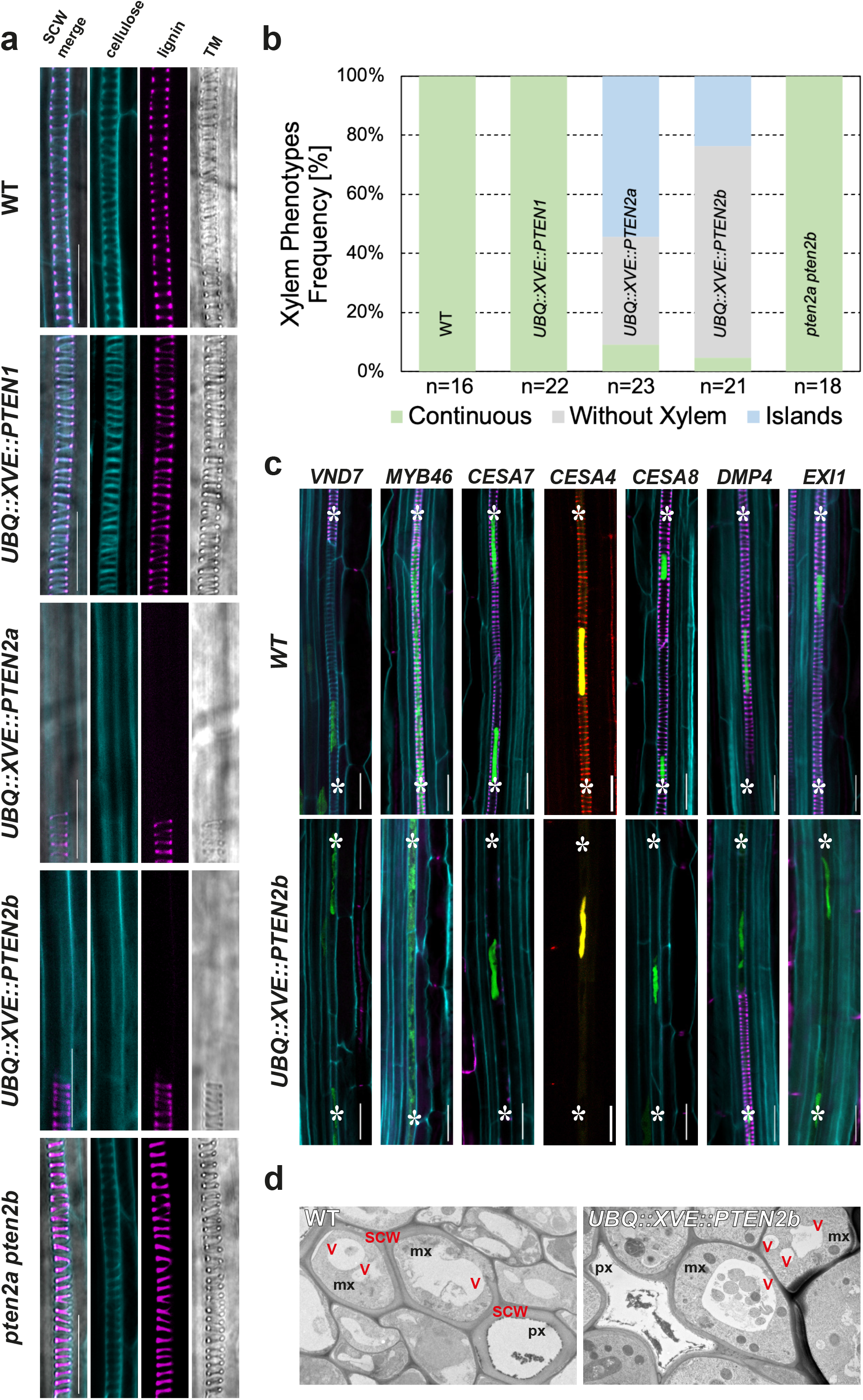
PTEN2s prevent xylem cell differentiation. **a**, Representative confocal images of protoxylem cells in the indicated genotypes stained with Calcofluor White (cellulose in cyan) and Fuchsin (lignin in magenta). Secondary cell wall (SCW) was also visualised by transmission light (TM). Scale bars represent 20μm. **b**, Quantification of xylem phenotypes observed in the roots displayed in a. The two phenotypes observed and scored were the total absence of xylem strands (without xylem) or the appearance of several protoxylem cells with SCW (islands). **c**, Representative images of indicated xylem differentiation markers in wildtype (WT) and seedlings incubated in 0.2μM estradiol for 48h to trigger *PTEN2b_ox_*. Roots were stained with Calcofluor White (cyan) and fuchsin (magenta). Marker lines: *VND7* (*VASCULAR RELATED NAC-DOMAIN PROTEIN 7*), *MYB46* (*MYB DOMAIN PROTEIN 46*), *CESA7* (*CELLULOSE SYNTHASE CATALYTIC SUBUNIT 7*), *CESA4* (*CELLULOSE SYNTHASE A4*), *CESA8* (*CELLULOSE SYNTHASE 8*), *DMP4* (*DUF679 DOMAIN MEMBRANE PROTEIN 4*), *EXI1* (*EXITUS 1*). Scale bars represent 20μm. **d**, Transmission electron microscopy images of differentiating proto- (px) and metaxylem (mx) cells in WT and *PTEN2b_ox_*. Xylem cells in WT formed thick secondary cell wall (SCW), vacuoles are enlarging in mx while px underwent programmed cell death. *PTEN2b* overexpression prevents SCW formation while mx cells contain multiple small vacuoles. Here px cell also underwent clearance.

Next, we assessed if xylem differentiation in *PTEN2b_ox_* was only delayed or lastingly inhibited by analyzing the expression of known xylem markers associated with xylem maturation. The expression of the protoxylem master regulator *VASCULAR RELATED NAC-DOMAIN PROTEIN 7* (*VND7*) and its downstream target gene *MYB DOMAIN PROTEIN 46* (*MYB46*) in seemingly non-differentiated xylem cells in *PTEN2b_ox_* validated that these cells committed to the xylem cell fate (Fig. 2c). Moreover, the expression of secondary cellulose synthetic machinery subunits as well as PCD-associated enzymes as a hallmark of xylem cell maturation can be detected in *PTEN2b_ox_* (Fig.2c and Extended Data Fig. 2a). These findings demonstrated that although the xylem cells reached their transcriptional maturity, they failed to lay down SCWs as confirmed by transmission electron microscopy (Fig. 2d). Notably, xylem cells failing to form SCWs contained vacuoles of altered morphology compared to wild type (Fig. 2d), implying correlation between the SCW formation and vacuolar morphology in xylem cells.

### PTEN2b overexpression modulates vacuolar and secretory vesicular trafficking of xylem cells

Despite its key role in protoxylem differentiation^27^, very little is known about xylem vacuolar biogenesis, mostly due to its relatively deep positioning within a root. Hence, we decided first to reassess vacuolar biogenesis in protoxylem developing cells by monitoring VHP1-GFP dynamics. Live-cell imaging, followed by 3D image reconstruction revealed the formation of elongated tubular structures in dividing meristematic cells (Fig. 3a). The subsequent enlargement of tubular compartments results from the fusion of small rounded vacuoles, giving rise to two large vacuolar compartments connected by a narrow tubular connection. *PTEN2s_ox_* effectively abolished the fusion of small vacuoles to the elongated tubular vacuole (Fig. 3b-c). Surprisingly, neither *PTEN2a_ox_* nor *PTEN2b_ox_* affected vacuolar morphology in root ground tissues (Fig. 3c). The latter suggested that xylem cells require a cell type specific vacuolar regulation. To corroborate the prevention of the xylem tubular vacuole enlargement and its globularization in *PTEN2sox*, we assessed the trafficking pathways summarized in Figure 4a. Similar to VHP1, VHA-a3 was present on the membranes of both tubular and small vacuoles in *PTEN2b_ox_* suggesting that the direct ER-to-vacuole trafficking pathway is functional (Fig. 4b). Next, we aimed to assess TGN-dependent vesicle delivery to the vacuole by analyzing an artificial cargo ToIM^28^ comprising a soluble vacuolar sorted part RFP_AFVY_ and GFP retained in the cytoplasm. Overexpression of *PTEN2b* precluded the loading of RFP_AFVY_ into the vacuole, as well as other cargos known to be transported to the vacuole (Fig. 4c-e and Extended Data Fig. 4a). Together, these observations indicated that *PTEN2b* regulates vesicle trafficking from TGN to the vacuole. This result was consistent with the previously reported PTEN2a localization at TGN^15^. Similarly, PTEN2b exhibits an early colocalization with membrane tracer FM4-64 and TGN marker VHA-a1 (Extended Data Fig. 4b-c). However, failed vacuolar delivery of these cargos cannot directly explain the inability of xylem cells to build a SCW. With the onset of SCW synthesis, the primary cell wall CELLULOSE SYNTHASE (CESA) enzyme complexes ceased to be delivered to the plasma membrane and are gradually removed^29^. As CESA6 is a primary cell wall CESA subunit, it may be plausible to hypothesize that the sequestering of this subunit in the vacuole is critical for the secondary cellulose synthase complex to assemble and/or be active. Although CESA6 colocalized with PTEN2b in discrete punctae, *PTEN2b* overexpression still abolished the xylem SCW formation in *cesa6* genetic background (Extended Data Fig. 4d-e), invalidating our hypothesis. SCW formation however also depends on the cellular secretion pathway as hemicellulose, lignin monomers and biosynthetic enzymes must be delivered to the apoplast before crosslinking^30–32^. For example, LACCASE 17 (LAC17) is an enzyme essential for lignin biosynthesis that in wild type plants can be found in the apoplast following the SCW spiral pattern^33^. Surprisingly, in *PTEN2b_ox_* LAC17 was not secreted, but rather retained inside the cell forming aggregates between the vacuoles (Fig. 4f-g), explaining the lack of lignin formation as a part of xylem SCW in *PTEN2b_ox_*. Although the vacuolar and secretion trafficking pathways may converge at multi-vesicular-bodies (MVBs), CESA6 and LAC17 aggregates created in *PTEN2b_ox_* did not always colocalize (Extended Data Fig. 4f). The latter suggests that PTEN2b can affect MVBs (as confirmed by Rha1 marker line) but also it may affect TGN downstream pathways at different levels (Extended Data Fig. 4g). In stronger *PTEN2b_ox_* we even occasionally noticed in epidermal cells RABG3f (RAB7 GTPase HOMOLOG) positive aggregates in a grape-like structures seemingly unable to fuse (Extended Data Fig. 4h), further supporting our findings in xylem tissue.

**Figure 3:**
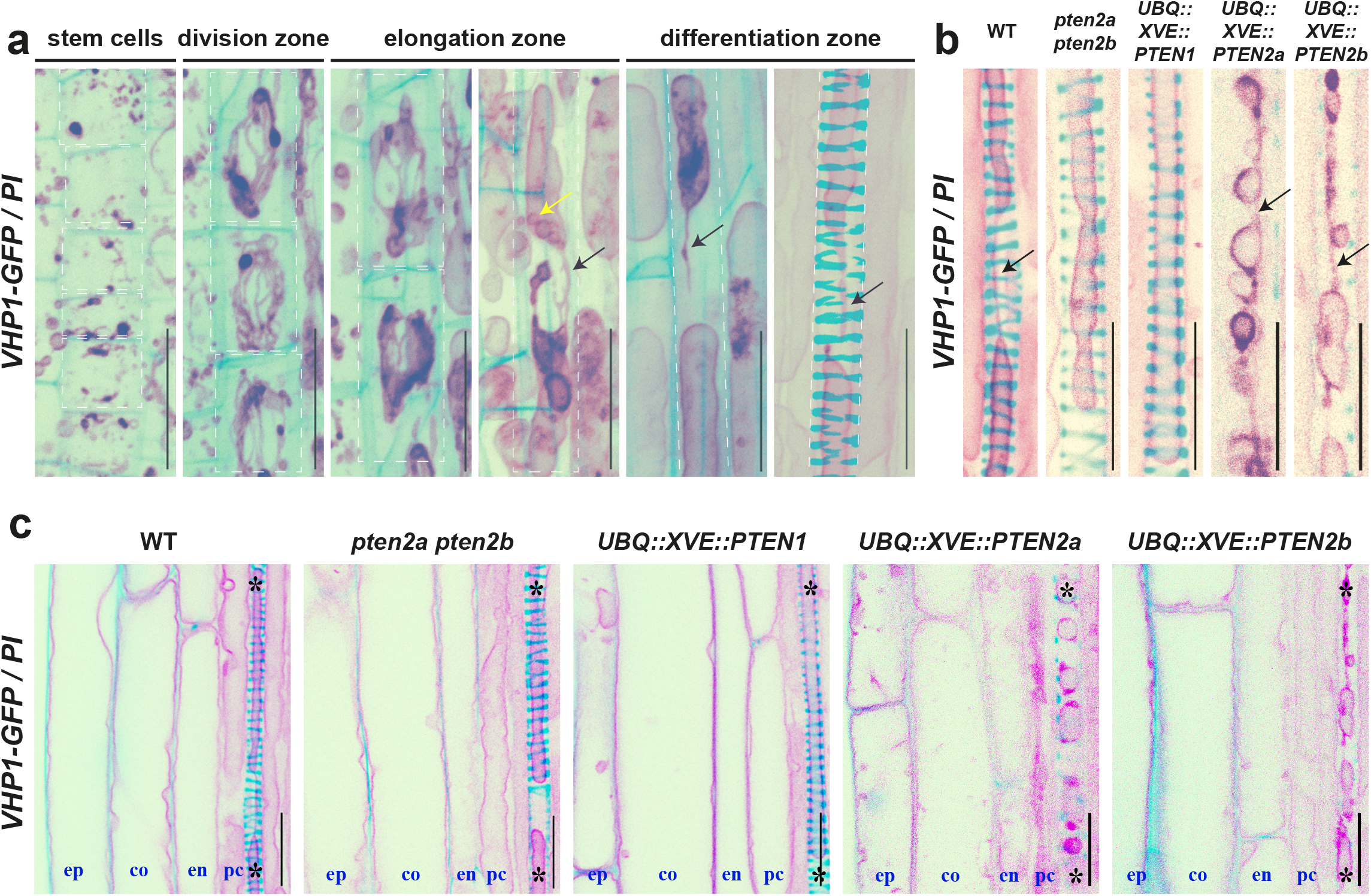
Regulation of xylem vacuolar biogenesis regulation by PTEN2s. **a**, 3D reconstruction of VHP1-GFP decorated vacuolar compartments in protoxylem cells in wild type plants at progressive developmental stages counterstained with propidium iodide (PI) to label cell wall. Yellow arrows mark small vacuole-like compartments and black arrows mark tubular connecting membranes. For easier visualization, protoxylem cells margins were squared by a white dashed line. Scale bars represent 20μm. **b**, Representative images of mature protoxylem cells in the indicated genotypes, visualized as in a. Scale bars represent 20μm. Black arrows mark tubular connecting membranes. **c,** Comparison of vacuolar morphology in mature epidermis (ep), cortex (co), endodermis (en), pericycle (pc) and protoxylem (asterisk) between indicated genotypes. Scale bars represent 20μm.

**Figure 4:**
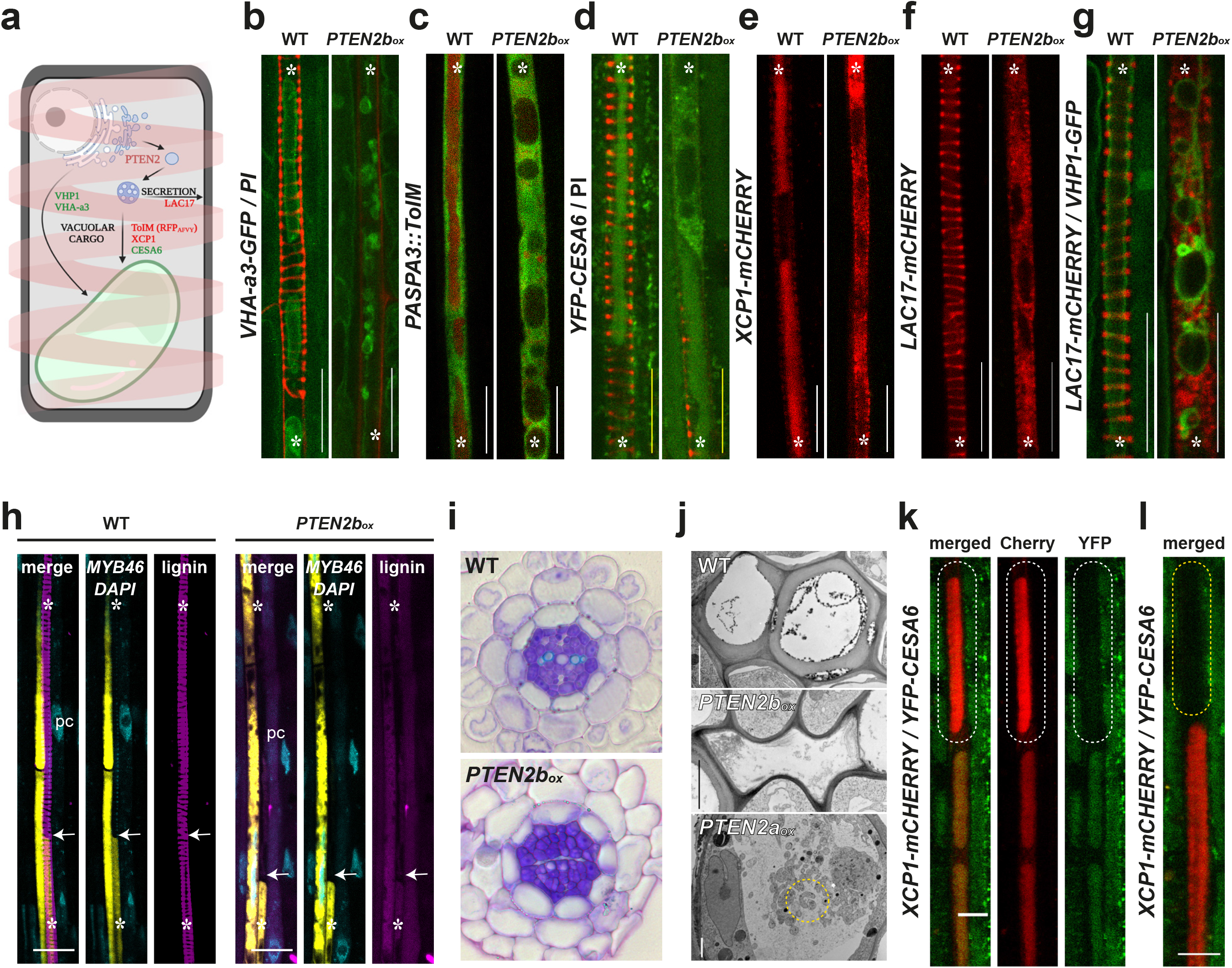
PTEN2s inhibit vacuolar and secretion trafficking pathway in xylem cells but not PCD. **a**, Schematic representation of analysed trafficking pathways important for xylem cell differentiation. **b-g**, Representative images of the corresponding xylem cells in wild type (WT) and seedlings with 0.2μM estradiol- mediated *PTEN2b* induction for 48h visualizing different trafficking markers: tonoplast marker VHA-a3 (VACUOLAR PROTON ATPASE A3) (**b**), xylem specific promoter *PASPA3* (*PUTATIVE ASPARTIC PROTEINASE A3*) driving expression of *ToIM* (tonoplast integrity marker) showing GFP in ER and a vacuolar targeted mRFP (**c**), vacuolar cargos CESA6 (CELLULOSE SYNTHASE SUBUNIT A6) (**d**) and XCP1 (XYLEM CYSTEINE PEPTIDASE 1) (**e**), secreted cargo LAC17 (LACCASE 17) (**f**). **g**, non-secreted LAC17 is not delivered into VHP1-labeled vacuoles. **h-j**, PCD execution occurs even without SCW formation in *PTEN2b_ox_*. **h**, DAPI-stained nuclei are absent in the cells where the xylem-specific expression of MYB46 ceased due to the PCD execution (white arrows). Scale bars represent 20μm. **i**, Toluidine-stained root cross- sections of the indicated genotypes. Xylem secondary cell wall stains bright blue as visible in WT but absent in *PTEN2b_ox_* overexpression where xylem cells appear collapsed as in transmission electron microscopy images (**j**). **j,** Transmission electron microscopy images of the indicated genotypes. Notice a high number of small vacuoles and aggregates in *PTEN2b_ox_*. Yellow dashed circle highlights a cup-shaped phagophore. Scale bars represent 2μm. **k**, Xylem vacuole acidification in wild type prior PCD. Note faster fading of pH-sensitive YFP in comparison to pH-tolerant mCHERRY. The cell where acidification occurs is encircled with a white dashed line. Scale bars represent 10μm. **l**, continuation cell from k, where PCD is executed and mCHERRY signal disappears too. Scale bars represent 10μm

Since vacuolar-driven PCD is the final step of xylem tissue maturation, we evaluated PCD execution in xylem cells upon *PTEN2b* overexpression. Remarkably, PCD execution can still be detected in cells uncapable of forming SCW, as manifested by the absence of organelles such as the nuclei (Fig. 4h). Xylem cells incapable to form SCW ultimately collapse, as seen in orthogonal sections stained with toluidine blue and transmission electron microscopy images (Fig. 4i-j).

Interestingly, a sharp quenching of YFP signal in comparison to mCHERRY could be detected prior to PCD execution (Fig. 4k-l). The different pH fluorescence of YFP and mCHERRY proteins^34^ may be accounted for this phenomenon, suggesting timed vacuolar acidification only in the last steps of xylem differentiation. The visibility of YFP fluorophore inside xylem vacuoles suggests that these vacuoles have a milder pH^35^, similar to storage vacuoles. Hence, this phenomenon may explain the cell-specificity of PTEN2s action on vacuoles in xylem tissues but not in other cell types (Fig.3c).

Together, our results showed that *PTEN2s* overexpression restricts tubular xylem vacuoles from enlarging. This phenotype is opposite to the phenomenon occurring during EV division, where even membrane tubularization is essential to create a furrow surrounding the incipient cleavage site (Extended Data Fig. 4i).

### *PTEN2* function was conserved through evolution before vascular plants appearance

Contrary to *PTEN2s*, overexpression of *PTEN1* did not affect vacuolar formation nor xylem differentiation (Figs. 2a-b and 3b-c). This observation raised the question whether the duplication of PTEN enzymes and their vacuolar remodeling function was an evolutionary prerequisite that contributed to the emergence of vascular plants (Fig. 5a). To answer this question, we applied a phylogenomic strategy for identifying orthologs in the green lineage. We blasted Arabidopsis *PTEN2b* full-length amino acid sequence against proteome assemblies of 142 plant species from chlorophytes to angiosperms (Fig. 5b, Supplementary Table 1). Detailed analysis revealed that PTEN enzymes from the green lineage can be divided into 3 subfamilies: algal PTEN, PTEN1 and PTEN2 (Fig. 5b). Algal PTEN subfamily includes *Chlamydomonas reinhardtii* PTEN (CrPTEN) that interestingly clusters with the referent human (HsPTEN) isoform. PTEN1 subfamily contains previously mentioned Arabidopsis PTEN1 (AtPTEN1), while PTEN2 subfamily clusters Arabidopsis isoforms (AtPTEN2a and AtPTEN2b) with *Marchantia polymorpha* PTEN2 isoform (MpPTEN2). The divergence of the *PTEN2* gene subfamily could be traced back to the origin of the *Phragmoplastophyta* clade (Fig. 5a) as confirmed by the absence of xylem and vacuolar phenotypes when overexpressing *CrPTEN* in Arabidopsis seedlings (Fig. 5c-d). Moreover, we observed that *PTEN2* genes conserved their functions even in the non-vascular plant *Marchantia* as the effect of overexpressing *MpPTEN2* mimicked *AtPTEN2b* overexpression. These observations suggest that vacuolar remodeling and xylem differentiation are *PTEN2*-specific functions that remained highly conserved across land plants despite hundreds of millions of years of evolution^36^.

**Figure 5:**
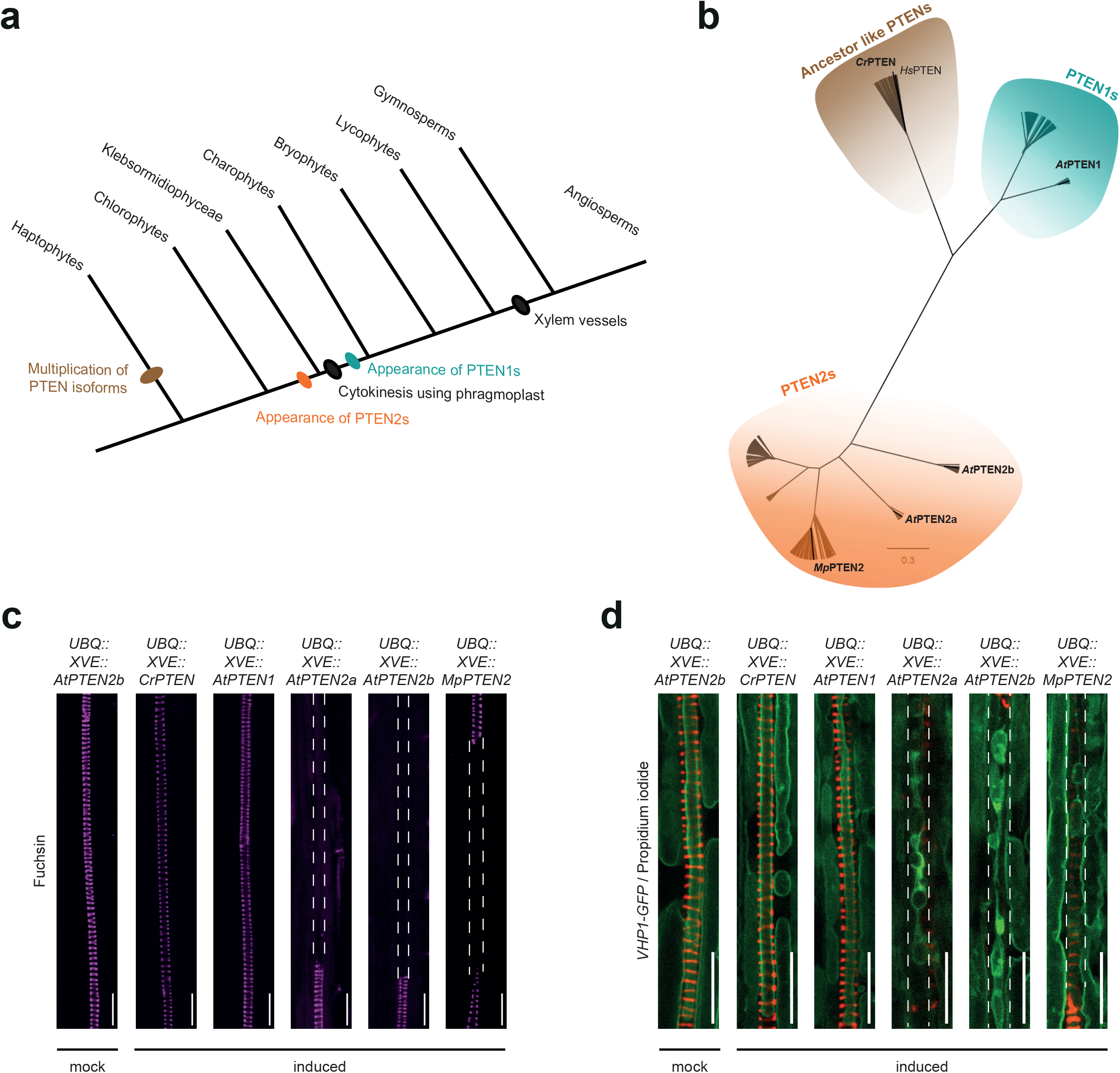
PTEN2s functions in vacuolar fusion and xylem differentiation were conserved through evolution. **a**, Schematic tree of the evolution of plant PTEN subfamilies. **b,** Phylogenetic tree of 418 plant PTENs from 142 plant species. For simplification, only the isoforms of species of interest have been represented. Details about all the sequences and the complete distribution of the isoforms in the three sub- families (ancestor-like PTENs, PTEN1s, PTEN2s) can be found in Supplementary Table 2. **c**, Representative confocal microscopy images of fuchsin-stained protoxylem strands from roots grown on mock conditions or upon 2µM estradiol-mediated induction for 48h of *Chlamydomonas reinharditi* PTEN (Cr*PTEN)*, *Arabidopsis thaliana PTEN1* (*AtPTEN1*), *PTEN2a* (*AtPTEN2a*), and *PTEN2b* (*AtPTEN2b*) and *Marchantia polymorpha PTEN2* (*MpPTEN2*). **d**, Representative confocal microscopy images of vacuolar morphology in mature xylem cells (VHP1-GFP labels tonoplast, propidium iodide stains cell wall) of inducible over-expressor lines of the different PTEN isoforms mentioned above. *PTEN* over-expression was induced by 2µM estradiol for 48h. Protoxylem gaps are highlighted with white dashed lines. Scale bars represent 20µm.

### PTEN2 function lies in its N-terminal domain that determines its subcellular localization

Cross-examination of PTEN sequences from the three subfamilies revealed extended N-terminal and C-terminal sequences in PTEN2s compared to other two subfamilies (Fig. 6a). To test whether these sequences may explain PTEN2s functional divergence from other PTEN subfamilies, we overexpressed N-terminal and C-terminal truncated versions of *At*PTEN2b (PTEN2b^ΔNter^ and PTEN2b^ΔCter^, respectively). Deletion of PTEN2b C-terminal sequence did not alter PTEN2b -dependent suppression of xylem continuity, whereas the lack of N-terminal sequence (*PTEN2b ΔNter*) inhibits this enzyme’s ability to impair xylem development (Fig. 6b) or vacuolar biogenesis (Fig. 6c). Furthermore, we were able to pinpoint the functional necessity of 57 AA of PTEN2 N-terminal domain (PTEN2b^Δ1-131^) upstream of its phosphatase catalytic domain (Fig. 6a-b). This conserved N-terminal part of PTEN2s makes it less hydrophobic in comparison to N-terminal of PTEN1, supporting the functional divergence between orthologues (Extended Data Fig. 6a-b). As expected, the N-terminals swapping from PTEN1 to PTEN2b (N1-PTEN2b) abolished PTEN2b function (Fig. 6b), confirming the functional specificity of PTEN2 N-terminus. Further *in silico* mining pointed out an enrichment in intrinsically disordered domains in the PTEN2 N-terminus suggesting the importance of macromolecular interaction partners in achieving stable PTEN2 three-dimensional structure (Extended Data Fig. 6c). Notably, the PTEN2b variants lacking complete (PTEN2b^Δnter^) or partial N-terminus (PTEN2b^Δ1-131^) could not properly localize to TGN (Fig. 6d and Extended Data Fig. 6d-f), where in addition to the cytosol PTEN2s normally accumulate^15^(Extended Data Fig 4c). PTEN1 isoform, shown to be restricted to pollen grains^37^, when expressed from the *PTEN2b* promoter cannot be detected in *PTEN2b* expressing tissues. Here it could be speculated that the long PTEN2 N-terminus (not present in PTEN1) in important not only for TGN localization but also for protein stabilization (Fig. 6d). Hence, it appears possible that PTEN2b Nter-mediated TGN anchoring is critical for PTEN2 functionality in vacuolar remodeling and contribute to the evolution of land plants

**Figure 6:**
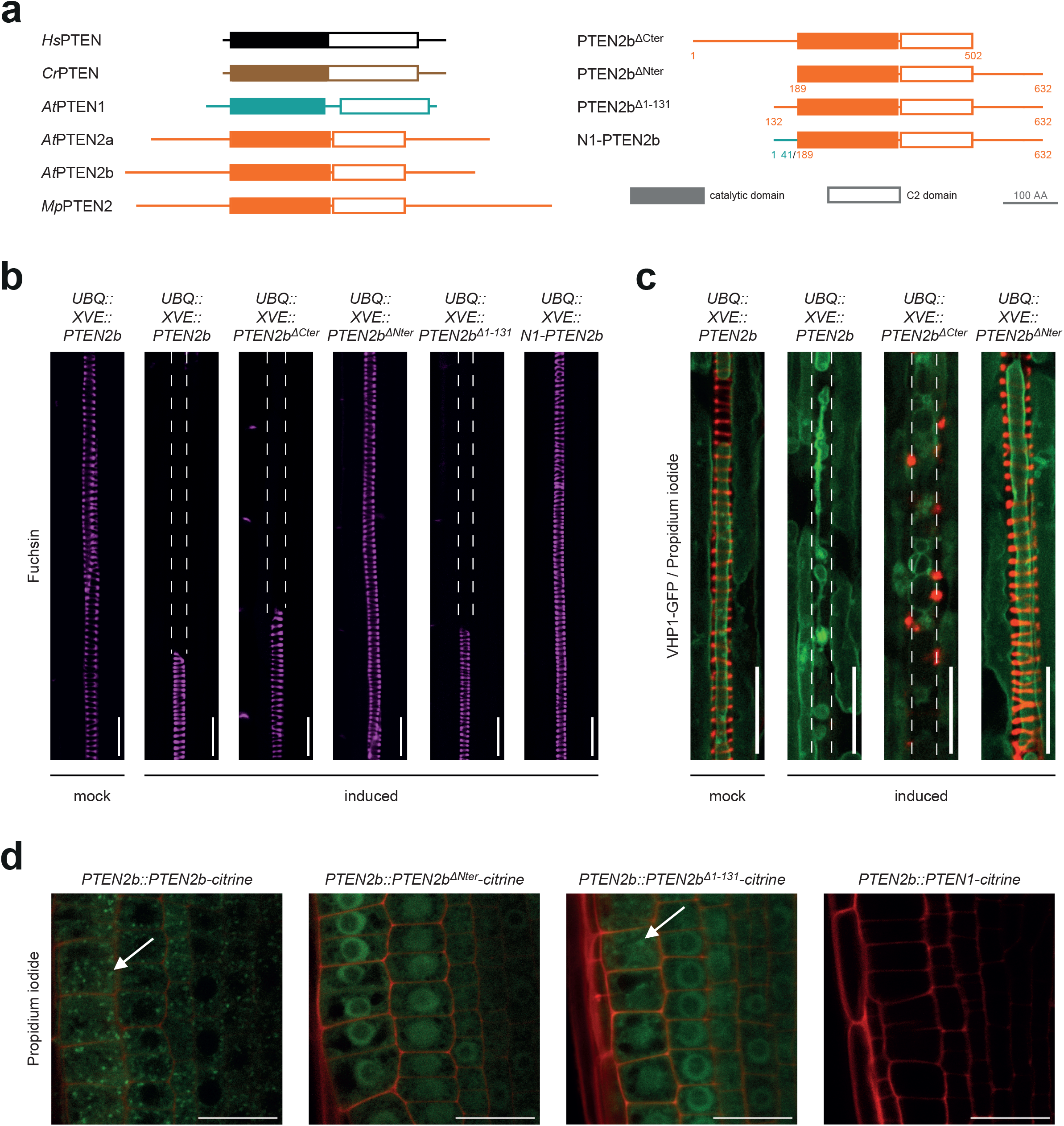
A conserved domain within PTEN2s N-terminal sequence is critical for their functionality and TGN anchoring. **a**, Schematic representation of the PTEN enzymes from *Homo Sapiens* (*Hs*), *Arabidopsis* (*At*), *Marchantia* (*Mp*) and *Chlamydomonas* (*Cr*) analysed in this study. On the right are represented the truncated version of PTEN2b without the entire C- (PTEN2b^ΔCter^) or N-terminal (PTEN2b^ΔNter^) sequences, PTEN2b with a partial N-terminal sequences (PTEN2b^Δ1-131^) and the hybrid version with PTEN1 N-terminal (N1-PTEN2b). Colour filled boxes represent phosphatase catalytic domains whereas empty squared boxes represent C2 domains. **b**, Representative confocal microscopy images of fuchsin-stained protoxylem strands after 2µM estradiol-mediated induction for 48h of indicated PTEN2b versions. Protoxylem gap cells are highlighted with white dashed lines. **c**, Representative images of vacuole morphology in mature xylem cell upon 48h overexpression of indicated PTEN2b variants. VHP1-GFP labels tonoplast, while propidium iodide labels cell wall. Protoxylem gap cells are highlighted with white dashed lines. **d,** Representative confocal images of 6-day-old plants harbouring indicated constructs illustrating the dependence of PTEN2b localization at TGN to its N-terminal. Scale bars represent 20µm.

## DISCUSSION

Eukaryotic cell compartmentalization occurred evolutionary concomitant with cell enlargement as the plasma membrane surface was not sufficient to provide all membrane-dependent functions^38^. There are different hypotheses explaining the origin of different membrane-bound organelles such as the endosymbiotic origin of mitochondria and chloroplasts, *de novo* formation of peroxisomes or transformation of existing endomembrane structures into new ones^39–41^. PSV formation represents an example when an existing compartment is remodeled into an organelle with a different function^22^. Yet, very little is known about the mechanisms underpinning this functional reprogramming process. Here we showed a PTEN2-mediated mechanisms by which numerous PSVs are formed by the fragmentation of EVs following a cleavage furrow-resembling mechanism (Fig. 1). Without PTEN2 enzymes the EV becomes hyper compartmentalized instead of fragmented. Interestingly, the observed EV membrane invaginations in *pten2a pten2b* mutants resemble formation of the mitochondrial cristae or chloroplast thylakoid membranes especially when the membrane is seen as pinched off (Fig. 1b)^38, 41, 42^.

As a typical cleavage furrow requires cytoskeleton involvement, it is tempting to speculate the importance of the cytoskeleton during vacuole division. Although actin was reported to be important in lytic vacuole tubularization, it is not clear if actin prevents vacuole expansion by physical constriction or by preventing actin-dependent membrane delivery to the vacuole^43^. Notably, vacuole invaginations can occur in a cytoskeleton independent fashion as reported during microautophagocytosis in yeast^44^. Moreover, the dynamics of contractile vacuoles present in protists depends rather on membrane tethering complexes than on the activity of cytoskeletal elements^45, 46^. Consistent with the coupled occurrence of fusion and fission events in membrane homeostasis, our work revealed a *PTEN2*-mediated effect on vesicle fusion in xylem cells (Fig. 4). Inducible overexpression of *PTEN2s* potentiated tubular vacuole structures by preventing small vacuoles to fuse with it (arrows in Fig. 3b). Vacuole tubularization (as the most extreme form of membrane bending) is actually a core phenomenon necessary to form a cleavage furrow-like structure during EV division (Extended Data Fig. 4i). In *pten2a pten2b* mutants, vacuolar fission is hindered by the failure to form a symmetric ring of tonoplast invaginations at the division site (Fig. 1b). Coincidentally or not, plant PTEN2s appeared exactly in the Phragmoplastophyta clade and diverged their function from ancestral PTENs with the loss of cleavage furrow as cytokinesis mechanism^47^.

*PTEN2s* overexpressing lines provided a critical genetic tool to research the cell type specificities of vesicle trafficking in a developmental context. In this study we focused on the cell trafficking during xylem cell differentiation into a water conducting unit. We showed that *PTEN2b_ox_* prevents SCW formation in xylem cells, partially by inhibiting LAC17 secretion to the apoplast. As mutations in hemicellulose biosynthetic genes translate into xylem phenotypes, a potential suppressed delivery of hemicellulose to the apoplast may explain the lack of SCW cellulose in xylem cells upon *PTEN2b_ox_* induction^48–50^. Furthermore, our current knowledge about xylem formation indicates that concomitant with SCW formation, hydrolytic enzymes necessary for PCD execution are loaded into the vacuole. Vacuoles store these enzymes in an inactive form until SCW formation is completed, ensuring the correct timing of PCD execution^51^. Our observations indicated that vacuolar acidification occurs just prior to PCD execution (Fig. 4k), suggesting a mechanism for the activation of hydrolytic enzymes. Subsequently, the vacuole swells, the tonoplast’s permeability changes and finally the vacuole collapses releasing its content into the cytoplasm^52, 53^. This process is thought to trigger a rapid cytoplasmic content degradation^54^. However, we showed that PCD-associated genes are correctly expressed and that protoxylem cells undergo cell death in the absence of SCW formation and without the formation of a large central vacuole, contrary to the expected sequence of xylem differentiation events^55^. Remarkably, previous studies reported autophagy as responsible for the gradual cellular content hydrolysis and reduced cytoplasmic density observed during the SCW biosynthesis^53, 56^. Thus, autophagy may be an alternative mechanism for xylem cell clearance when vacuole-mediated pathway is inhibited. Indeed, we observed the creation of multiple cup-shaped vesicular structures resembling phagophore upon high *PTEN2a* induction (Fig. 4d) as well as that RabG3f positive small vacuoles creating grape-like aggregates (Extended Data Fig. 4h). It has been reported that another member of the same RabG3 subfamily, RabG3b can either stimulate or inhibit both autophagy and xylem formation, depending on its activation status^57^. Interestingly, the autophagy resembling pathway occurs also when massive amounts of synthesized proteins have to be delivered to the vacuole, as it occurs for seed storage proteins^58, 59^. Similarly, some proteases necessary to mobilize the PSV content during germination was shown to also skip Golgi/TGN and directly from ER translocate to the vacuole^60, 61^. Utilization of this direct ER-to-vacuole pathway may explain the absence of germination defects in *pten2a pten2b* seedlings incapable to form conventional PSVs. This notion is supported by our result that the direct ER-to-vacuole trafficking route remains unaffected in *PTEN2b_ox_*, evident by the presence of VHP1 and VHA-a3 in xylem vacuolar membranes (Fig. 3b and 4b)^18^. Further investigation is needed to elucidate the exact downstream players in PTEN2s signaling cascade, and distinguish the biological importance of its dual phosphatase activity^10^, especially in the new light of its evolutionarily novel N-terminal. However, it is tempting to speculate that the *PTEN2* gene family diverged in green lineage to control vacuolar morphology and dynamics as their emergence coincided with vacuole enlargement during evolution.

## METHODS

### Plant materials and growth conditions

*Arabidopsis thaliana* ecotype *Columbia-0* (*Col-0*) was used as wild-type control in all cases. Seeds of *pten2a (*SALK_114721)*, pten2b* (SALK_120020) and *cesa6* (SALK_004587) were obtained from the Nottingham Arabidopsis Stock Centre and combined by crossing. Homozygous lines were selected by genotyping using the primers listed in Supplementary Table 2. The following transgenic lines used in this study were described elsewhere: *MYB46::GFP^62^, DMP4::H2A-GFP^63^, EXI1::H2A- GFP^63^, PASPA3::H2A-GFP^63^, RNS3::H2A-GFP^63^, SCPL48::H2A-GFP^63^, PASPA3::ToIM^28^, VHA-a3::VHA-a3-GFP^18^, VHP1::VHP1-GFP^64^, VHA-a1::VHA-a1-RFP^65^, CESA6::YFP-CESA6^66^, LAC17::LAC17-mCHERRY^33^, BRI1::BRI1-mCITRINE^67^, UBQ::Rha1-YFP (W7Y)^68^*, *UBQ::RabG3f-YFP (W5Y)^68^*. These lines were combined with mutants or other transgenics by crossing. Following constructs (detailed information in Supplementary Table 3) were generated in this study: *TIP3;2::TIP3;2-GFP*, *UBQ::XVE::PTEN1*, *UBQ::XVE::PTEN2a*, *UBQ::XVE::PTEN2b*, *VND7::NLS-3xVENUS*, *CESA7::NLS-3xVENUS*, *CESA4::NLS-3xVENUS*, *CESA8::NLS-3xVENUS*, *XCP1::XCP1-mCHERRY*, *BFN1::NLS-dtTOMATO*, *PTEN2b::PTEN2b-CITRINE*, *PTEN2b::PTEN2b-mCHERRY*, *UBQ::XVE::CrPTEN*, *UBQ::XVE::MpPTEN2*, *UBQ::XVE::PTEN2b^ΔCter^, UBQ::XVE::PTEN2b^ΔNter^, UBQ::XVE::PTEN2b^Δ1-131^ UBQ::XVE::N1-PTEN2b, PTEN2b::* PTEN2b*^ΔNte^*- CITRINE, *PTEN2b:: PTEN2b^Δ1-131^ -CITRINE*, *PTEN2b::PTEN1-CITRINE*. Constructs were transformed into Col-0, *pten2a pten2b* and marker lines (unless indicated) using *Agrobacterium*-mediated floral dip transformation according to standard procedures. For *in vitro* growth, seeds were surfaced sterilized, stratified 2 days at 4°C and grown on 0.5 x Murashige and Skoog (MS, Duchefa) medium with MES buffer, pH 5.7, 0.7% agar, and 1% sucrose. Seedlings were grown in vertical plates under continuous light conditions. The estradiol inducible lines were either germinated or transferred to identical media (48h treatments) containing 0.2 µM or 2µM estradiol (ES, Sigma Aldrich). Seedlings were analyzed six days after germination unless specified otherwise.

### Cloning procedures

To generate *pPROMOTER::NLS-3xVENUS* reporter lines, the genomic region upstream the ATG of *VND7* (1596bp), *CESA7* (1153bp), *CESA4* (1939bp), *CESA8* (1949bp), or BFN1 (1975bp) was PCR-amplified using the primers listed in Supplementary Table 2. The resulting fragments were cloned into *pDONRPr-P1r* (Gateway) and subsequently recombined together with a *pENzeo-NLS-3xVENUS*^69^ plasmid into *EDO097pFR7m24G*^70^. Entry clone with *BFN1* promoter was recombined together with *pEN-L1-NLS-tdTOMATO-L2* (Gateway) into destination vector pK7m24GW2 (Gateway) plasmid following the manufacturer instructions (Gateway, Invitrogen).

*TIP3;2::TIP3;2-GFP* line was generated by synthetizing *TIP3;2* codding genomic sequence together with 845bp of promoter region upstream of ATG (as found in The Arabidopsis Information Resource Platform) and cloned into pDONRP4-P1r. Obtained entry clone was recombined with *pEN-L1-GFP-L2* into *EDO097pFR7m24G*^70^.

Protein overexpression was achieved by estradiol XVE system (Gateway plasmid pMDC7^71^). Coding sequences of *PTEN1*, *PTEN2a* and *PTEN2b* were amplified from Arabidopsis genomic DNA. *MpPTEN2* (Mapoly0016s0179) was PCR amplified from Marchantia cDNA. *CrPTEN* (Cre06.g308400) was in vitro synthesized (Invitrogen). DNA amplicons containing attB1-B2 sites were and recombined into *pMDC7* via *pEN207* (*PTEN1* and *PTEN2b*) or *pEN221* (*MpPTEN2* and *CrPTEN*) Gateway plasmids. *PTEN2a* was firstly cloned into *p17ACCD2P*, a plasmid created in this study. *p17ACCD2P* contains multicloning restriction sites: 5’- GAA TTC GAA GCT CGG TAC CCG GGG ATC CTC TAG AGT CGA CCT GCA GGC CCA TGG TGA CTA GTC AAG CTT – 3’ between attL1 and attL2 Gateway recombination sites, thus providing direct creation of an entry clone without BP reaction.

Translational reporters were created by amplifying promoter regions of: *PTEN2a* (1116bp), *PTEN2b* (1173bp) and *XCP1* (1601bp) and cloned into *pDONRPr-P1r*. Codding regions were amplified from whole seedling cDNA (for *PTEN2a*, *PTEN2b* and *XCP1*) or pollen enriched cDNA (*PTEN1*). Entry clones were made using pDONR207 except for *PTEN2a*, were *p17ACCD2P* was used. Final constructs were generated by recombining the entry clones into *pH7m34GW*. Similarly, the distinct *PTEN2b* protein variants (*PTEN2b^ΔNter^*, *PTEN2b^ΔCter^*, *PTEN2b^Δ1-131^* and *N1-PTEN2b*) were cloned in frame with CITRINE by LR recombination into *pH7m34GW* or for overexpression into *pMDC7*. Primers using for cloning can be found in Supplementary Table 2.

### Histological analysis

PSV biogenesis was visualized in epidermal cells of Arabidopsis embryos extracted from green siliques (stages I-III), yellow siliques (stages IV-V), dry seeds after 4h imbibition in water (stage VI) or 24-48h after imbibition in constant light conditions (stages VII-VIII). Cellulose (by Calcofluor White from Sigma-Aldrich) and lignin staining (Fuchsin from Sigma-Aldrich) were performed after seedlings clearing following ClearSee protocol as previously described^72^. DAPI (4′,6-diamidino-2-phenylindole, Sigma-Aldrich) staining used to verify PCD status of xylem cells was performed after fuchsin staining when seedlings were exposed to 50 μg/ml DAPI in 1x PBS (phosphate-buffered saline) with 1% TRITON-X for 1h with gentle shaking. After thorough washes in 1x PBS, seedlings were visualized with a confocal microscope. Live imaging of green fluorophores was performed upon propidium iodide (Sigma-Aldrich) or FM4-64 (Invitrogen) staining according to standard procedures^65^. Chemical treatment with Brefeldin A (BFA, Sigma-Aldrich) was performed for duration of 90 minutes in concentration of 50 μM in liquid 0.5 MS media. Transverse plastic sections of roots were performed and visualized as previously described^73^.

### Confocal microscopy and image analysis

Confocal laser-scanning microscopy images were obtained using either a Leica SP8 (in Fig. 4, Fig. 5 [b-g, k], Fig. 6d, Fig. 7c-d and Extended Data Fig. 5 [a, f-h]) or Zeiss LSM 780 microscopes. Blue dyes such as Calcofluor White and DAPI were excited at 405nm and detected at 430-485nm as well as PSV autofluorescence. Green fluorophores (GFP, CITRINE, Venus, YFP) were excited at 488nm and detected between 500-550nm. Red fluorophores and dyes (RFP, mCHERRY, tdTOMATO, propidium iodide, FM4-64 and fuchsin) were excited at 561nm and detected at 590-650nm. 63x Oil Plan-Apochromat DIC M27 objective was used to visualize Arabidopsis embryos, otherwise the 40x water LD C-Apochromat M27 objective was used on Zeiss LSM 780 microscope. For two photon fluorescence excitation, Mai Tai XF (Spectra-Physics) laser at 980nm was used to excite GFP, YFP and CITRINE, while InSight DeepSee (Spectra-Physics) at 1060nm was used to excite RFP and mCHERRY fluorophores. Here, 40x water HC PL IRAPO objective was used. Signal detection was collected with external detectors based on FITC (525/50nm) / TexasRed (617/70nm) filters. Images were processed in ImageJ or Imaris image processing software. When image colors were inverted and/or adjusted, all images belonging to one experiment were processed simultaneously. Scale bars were added in ImageJ.

### Transmission electron microscopy

The root of 7-day-old seedlings was mounted in 1-hexedecene (Sigma) on a carrier with a 2 mm diameter and high-pressure frozen using the Leica EM HPM100. Then the samples were substituted in 1% OsO_4_ for 6h at -90°C, followed by 3h at -60°C, 3h at -30°C, and 1h at 0°C. After one hour incubation, samples were rinsed twice with anhydrous acetone and incubated for 2h in 33% Epon/Araldite (Epon 812-Sigma, Durcupam ACM-Sigma, Dibutylphtalat-Sigma) in anhydrous acetone at 4°C. Next, samples were incubated in 66% Epon/Araldite in anhydrous acetone at 4°C overnight, and finally embedded in 100% Epon/Araldite. Samples were then trimmed using a glass knife and 70nm sections were cut with a diamond knife (DiATOME) using a ultramicrotome (Leica Ultracut UCT). Sections were assembled on a grid (2mmx1mm slit diaphragm, PLANO), coated with formvar (0.85% formvar in 1,2-dichlorethane). Contrast of the samples on the grids were enhanced with lead citrate. Samples were examined using the FEI Tecnai G2 Spirit transmission electron microscope with two digital CCD cameras (Gatan Orius 1000, FEI Eagle).

### RNA-extraction, RT-qPCR and Western Blot analysis

Total RNA was extracted from 7-day-old seedlings using RNeasy® Plant Mini-Kit (QIAGEN) and treated with RNase-Free DNase (QIAGEN). cDNA synthesis was performed using RevertAid First Strand cDNA synthesis kit (Thermoscientific). RT-qPCR was performed using KAPA SYBR® FAST (KAPA BIOSYSTEMS), primers listed in Table S1 and 2µL of 1:10 dilution of cDNA. All reactions were performed in triplicates. Expression data derived from Cp values calculated according to the second derivative maximum method (LightCycler®LC480 II, Roche) and normalized to the expression of *PDF2* (At4g04890). To detect PTEN2-CITRINE protein in the transgenic lines, total proteins of 7-day-old seedlings were extracted using Laemmli buffer (v/v), separated in 12.5% (w/v) sodium dodecyl sulfate-polyacrylamide gel electrophoresis (SDS-PAGE) and transferred to an AmershamTM HybondTM 0.45µm polyvinylidene difluoride (PVDF) membrane (Merck). CITRINE-fusion proteins were detected with anti-GFP antibody (JL-8, Takara Bio Clontech, dilution: 1:5000, overnight incubation) and anti-Mouse IgG (Fc-specific)-Peroxydase (Sigma, dilution: 1/10000, 1.5hr incubation). Anti-Hsp90 (Agrisera, dilution: 1:2500, overnight incubation) and anti-Rabbit IgG (whole molecule)–HRP (Sigma A0545, dilution: 1:5000, 1.5hr incubation) were used as loading control. Chemiluminescence was revealed using Roti®-Lumin (ROTH) as substrate and imaged with a ChemiDoc Touch Imaging System (Bio-Rad).

### Phylogenetic analysis

Protein sequences of PTEN2b homologous from 142 plant species from Phytozome, National Center from Biotechnology Information and Plaza 4.0 databases were analyzed using the following criteria: E value cut off 10^-10^ (Phytozome), total score cut off 100 (NCBI) and score cut off 100 (Plaza). Next, we removed manually the sequences whose identity was higher than 99% within the same species as well as the incomplete sequences. We only represented one splice variant for each locus and remove miss-aligned sequences. The resulting sequences were aligned using CLUSTAL OMEGA algorithm and the tree was generated by using FigTree version 1.4.3 software (http://tree.bio.edu.ac.uk/software/figtree/) and color-coded edited manually.

### Bioinformatic analysis of physicochemical protein properties

The coding sequences of the N-terminal domain of *CrPTEN, AtPTEN1, AtPTEN2a, AtPTEN2b* and *MpPTEN2* were aligned (CLUSTAL OMEGA) and the presence of membrane binding domains was predicted using a BH score above 0.6 as described by Brzeska et al^74^. By using IUPred2A score^75, 76^, domains with values above 0.5 were assigned as highly probably intrinsic disorder domains.

## ACKNOWLEDGEMENTS

The authors thank Dr. K. Schumacher, Dr. Y. Oda and Dr. S. Fujita for kindly providing transgenic material, Dr. J. Westermann for providing us with Marchantia cDNA, Dr. A. Ruiz-Sola, Dr. J Alassimone and K. Kirchoff for great technical assistance. We thank J. Kusch and T. Scwartz from ScopeM for their support in handling the confocal microscopes and image processing and to University of Zürich TEM services for technical assistance. B.G. was financially supported by Vontobel, A.H. and T.M.D.C. by ETH-Foundation grants, and C.C., N.B-T., and E.R. by ETH core funding. This work was also funded by the Swiss National Foundation (SNF_31003A_160201 to A.R.-V.).

---

**Extended Data Figure 1:**
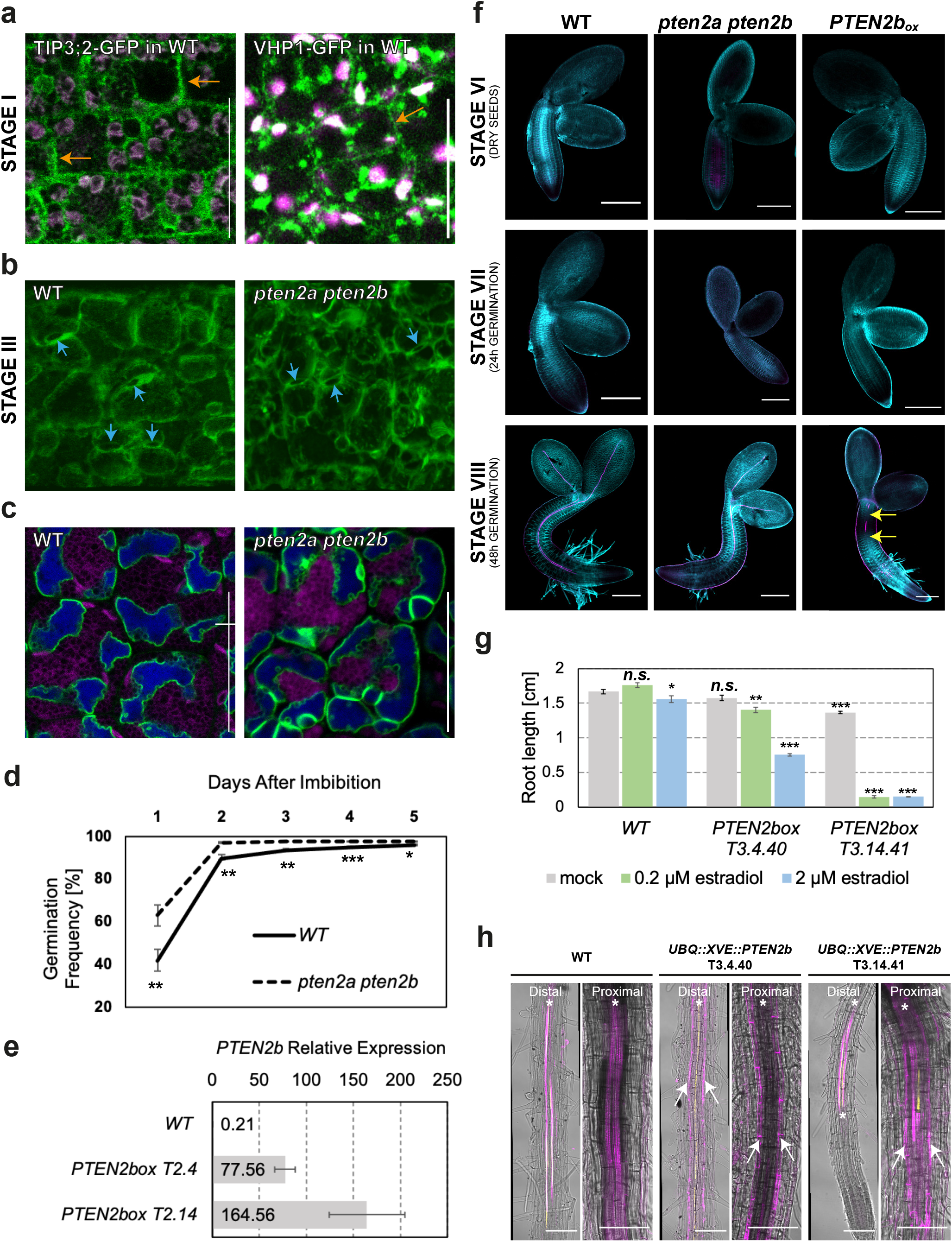
Aberrant EV division does not affect post-embryonic development. **a-c**, Representative confocal images of aquaporin *TIP3;2-GFP* (green) distribution during embryogenesis. **a**, Similar to TIP3;2, VHP1 tonoplast marker also labels ER. Magenta shows autofluorescence from chloroplasts visible in embryos isolated from green siliques. **b**, 3D maximal projection of a stage III vacuole corresponding to 2D image shown in Figure 1a. **c**, Small vacuoles labelled with TIP3;2 close to EV tonoplast preceding the cleavage furrow division. Magenta shows autofluorescence detected in red part of the spectrum. Scale bars represent 20µm. **d**, Germination rate between indicated genotypes. Error bars represent SE. n>500. **e**, Analysis of normalized, relative *PTEN2b* overexpression by qRT-PCR in two independent transgenic lines induced with 2μM estradiol for the duration of 6 days. Error bars represent SE among three independent biological replicates. **f**, Representative confocal images of embryos dissected from dry seeds, 24h and 48h after imbibition of the indicated genotypes, stained with Calcofluor White (cellulose in cyan) and fuchsin (lignin in magenta). Yellow arrows mark xylem discontinuities in seedlings with induced *PTEN2b* overexpression from imbibition (2μM estradiol). Note the appearance of differentiated xylem cells (magenta) only 48h after germination. Scale bars 200μm. **g**, Root length quantification of 6-days-old seedlings illustrate dose dependant effect of *PTEN2b* overexpression after 48h of estradiol induction. Error bars represent SE. n>40 **h**, *PTEN2b* overexpression prevents xylem differentiation in T3.4.40 line in both proximal and distal part of the root. Higher overexpression in the line T3.13.41 dramatically shortens the root length (**g**) but does not prevent xylem differentiation in younger distal root parts. Undifferentiated xylem is only labelled with MYB46 marker in yellow, while differentiated xylem is labelled with fuchsin staining for lignin in magenta, or white (overlap between yellow and magenta). Asterisk labels xylem position within vascular cylinder. White arrows label ectopic lignification in endodermis. Scale bars represent 100μm. n.s., not significant; ∗p<0.05; ∗∗p < 0.01; ∗∗∗p < 0.001.

**Extended Data Figure 2:**
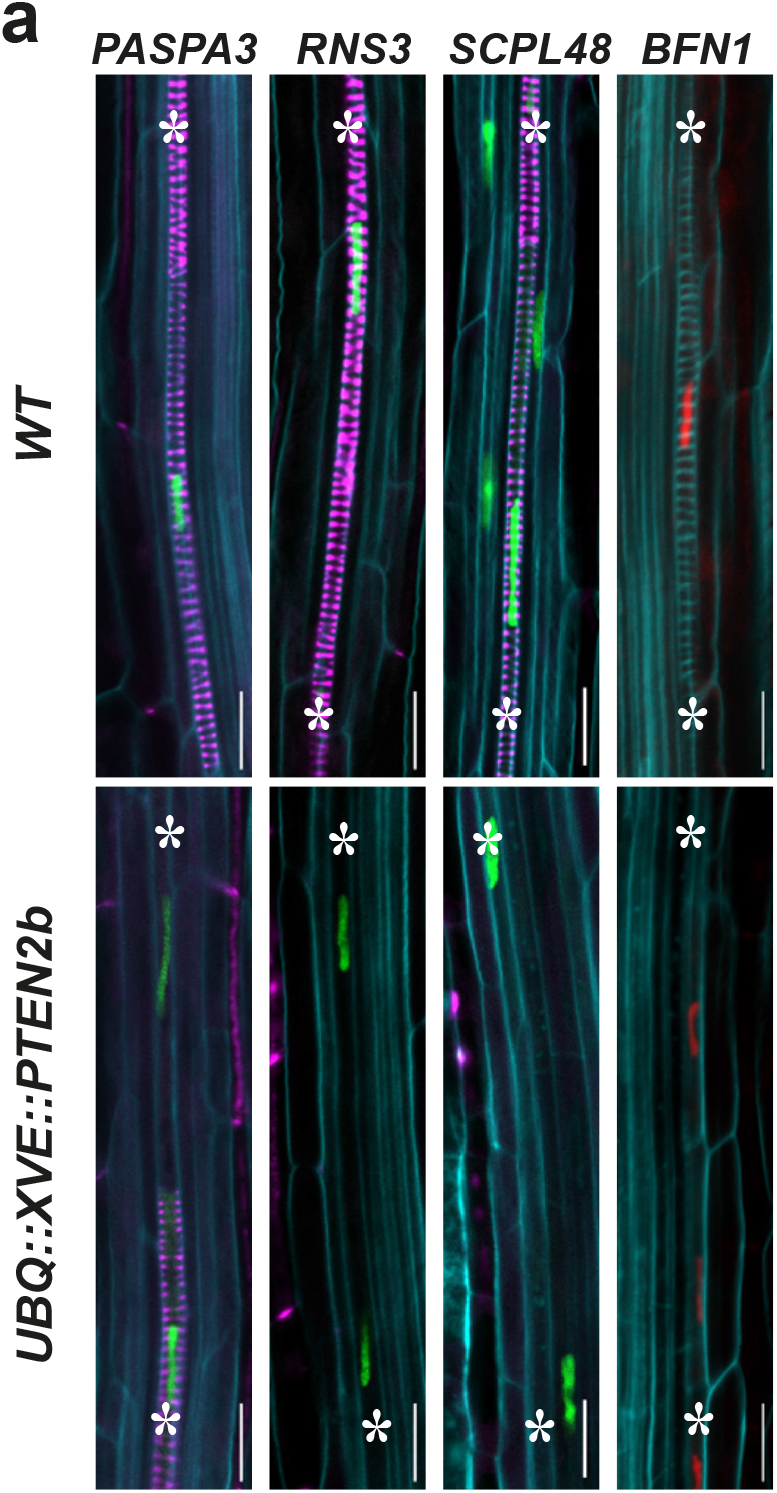
*PTEN2b* overexpression does not alter the expression of genes associated with xylem PCD execution. **a**, Representative confocal microscopy images of the mature protoxylem cells stained with Calcofluor White for cellulose (cyan) and fuchsin for lignin (magenta). *PTEN2b* was induced for 48h with 0.2μM estradiol. Note the expression of genes associated with PCD such as the *PUTATIVE ASPARTIC PROTEINASE A3* (PASPA3), *RIBONUCLEASE 3* (*RNS3)*, *SERINE CARBOXYPEPTIDASE-LIKE 48 (SCPL48)* and *BIFUNCTIONAL NUCLEASE 1 (BFN1)* can still be detected in seedlings with *PTEN2b* upregulation. Asterisks mark protoxylem strands. Scale bars represent 20μm.

**Extended Data Figure 4:**
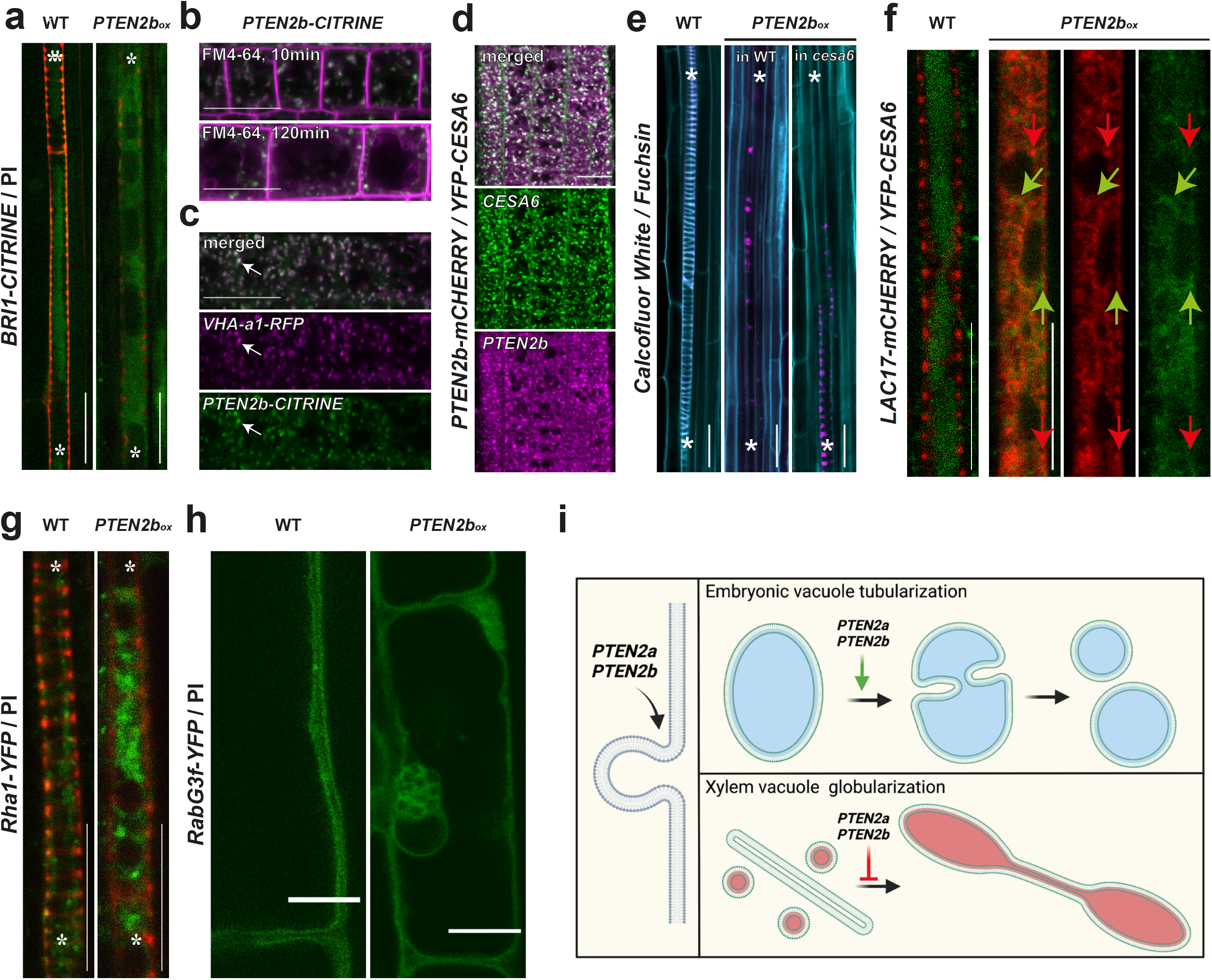
PTEN2b colocalize to TGN and impinges on vacuolar and cell secretion pathways. **a**, Brassinosteroid receptor BRI1 (BRASSINOSTEROID INSENSITIVE 1) cannot be delivered to xylem vacuoles upon *PTEN2b* overexpression. Seedlings were counterstained with propidium iodide (PI). **b**, PTEN2b colocalize with cellular compartments early labelled with FM4-64 (magenta). **c**, PTEN2b partially colocalize with PROTON ATPASE A1 (VHA-a1) in TGN (arrows). **d**, PTEN2b partially colocalize with CELLULOSE SYNTHASE SUBUNIT A6 (CESA6). **e**, *cesa6* mutant cannot rescue secondary cellulose building upon *PTEN2b* overexpression. **f**, Aggregates of vesicles carrying vacuolar destined cargo (CESA6 in green) and secretion cargo (LAC17 in red) do not colocalize. Arrows’ color corresponds to fluorophores and points the aggregates where proteins do not colocalize **g**, Multivesicular body (MVB) marker Rha1 (ARABIDOPSIS RAB HOMOLOG F2A) creates aggregates in xylem cells upon PTEN2b upregulation. **h**, Prevacuolar compartment and tonoplast marker RabG3f (RAB GTPASE HOMOLOG G3F) upon prolonged *PTEN2b* overexpression creates grape like structures in vicinity of the central vacuole in mature epidermal cells. Asterisk labels xylem strands. Scale bars represent 20μm. i, Schematic representation of membrane phenomena regulated by PTEN2s.

**Extended Data Figure 6:**
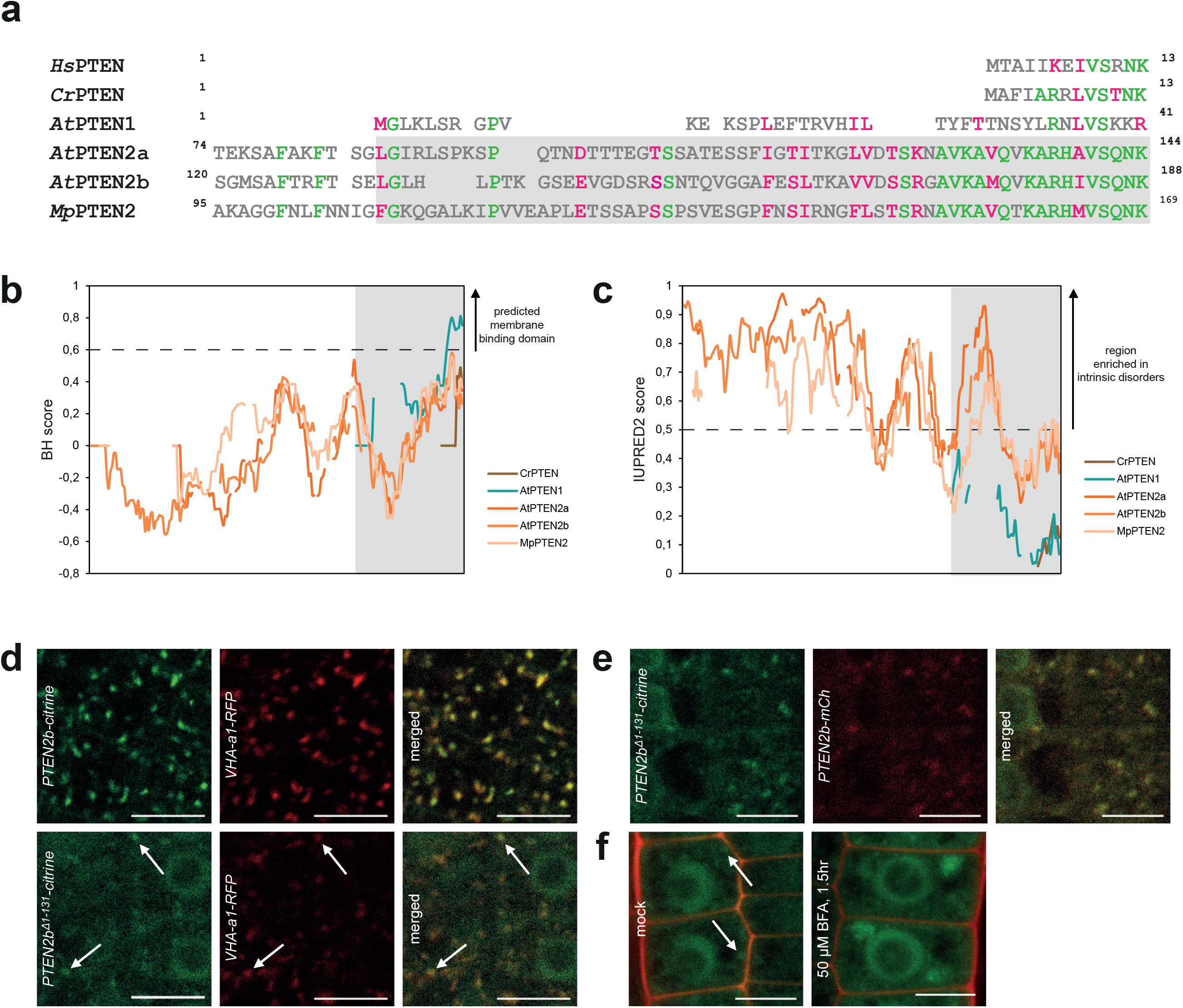
The N-terminal domains of PTEN1 and PTEN2s exhibit different biochemical properties that determines their subcellular localization. **a**, Alignment of N-terminal PTEN sequences from: human (*HsPTEN*), Chlamydomonas (*CrPTEN)*, Arabidopsis (*AtPTEN1, AtPTEN2a* and *AtPTEN2b), and Marchantia* (*MpPTEN2)* proteome assemblies obtained using CLUSTAL OMEGA. Identical amino acids are represented in green while similar amino acids are represented in magenta. **b**, Prediction of membrane binding domain in PTEN N-terminal sequences of indicated isoform using BH score^74^. Domains with values above 0.6 are predicted to be membrane binding domain. **c**, Prediction of intrinsic disordered region in N-terminal sequences of indicated isoform using IUPred2A score^75, 76^. Regions with values above 0.5 are supposed to be enriched in intrinsic disorders. The grey areas highlight the domain identified in PTEN2b (amino acid 132-188) as responsible for its functionality. **d**, Representative confocal images of 6 day-old plants expressing the TGN marker *VHAa1-RFP* together with *PTEN2b::PTEN2b-citrine* or *PTEN2b::PTEN2b^Δ1-131^-CITRINE*. Arrows indicate the position of some of the dotted structures observed for *PTEN2b::PTEN2b^Δ1-131^-CITRINE*. **e**, Representative confocal images of 6 day-old plants expressing *PTEN2b::PTEN2b^Δ1-131^-CITRINE* with *PTEN2b::PTEN2b-mCHERRY.* Please note the co-localization of both constructs. **f**, Confocal images of 6 day-old plants expressing *PTEN2b::PTEN2b^Δ1-131^-citrine* and treated with DMSO or 50 µM BFA for 1.5h. Scale bars represent 10µm.

**Supplementary Figure 1:**
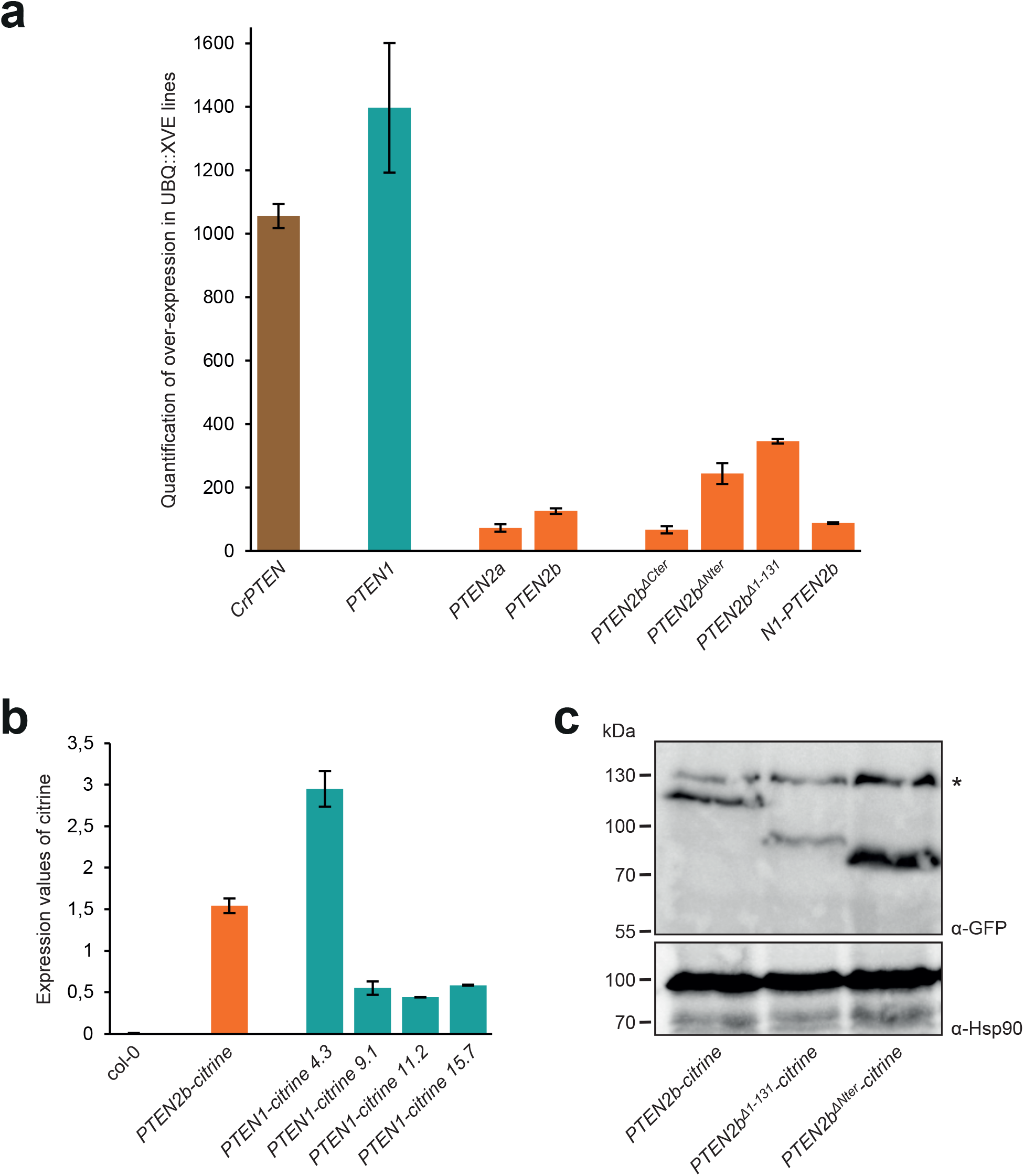
Transgenic lines validation (Supporting data for Figures 6 and 7). **a**, qPCR analyses confirmed the over-expression of PTEN in different inducible lines described in Fig. 6 and Fig. 7. RNA was extracted from roots of 7day-old plants treated with DMSO or 2µM estradiol for 48hrs. Expression values of the different genes of interest in estradiol-treated plants were normalized by the corresponding expression values measured in DMSO-treated plants. Values represent the mean of two biological replicates (both including three technical replicates), error bars indicate the standard deviation. **b,** qPCR analyses revealed the presence of *CITRINE* mRNA in independent *PTEN2b::PTEN1-CITRINE* that do not exhibit any fluorescence signal in root cells (Fig 6). Values represent the mean of three technical replicates, error bars indicate the standard deviation. **c,** The expression of the different Nter-truncated versions of PTEN2b tagged with citrine (Fig 6) was confirmed by Western blot using anti-GFP antibody. Anti-Hsp90 was used as a loading control. The star indicates the presence of an unspecific band.

**Supplementary Table 1:** Protein sequences used to build the phylogenetic tree.

| Organism | Protein name | Order | Clade | Database |
| --- | --- | --- | --- | --- |
| *Acorus americanus* | Acora.04G145300 | Acorales | Monocotyledones | phytozome |
| *Acorus americanus* | Acora.11G167200 | Acorales | Monocotyledones | phytozome |
| *Aegilops tauschii* | LOC109747383 | Poales | Monocotyledones | NCBI |
| *Alyssum linifolium* | Alyli.0051s0214 | Brassicales | Eudicotyledones | phytozome |
| *Alyssum linifolium* | Alyli.0020s0220 | Brassicales | Eudicotyledones | phytozome |
| *Alyssum linifolium* | Alyli.0204s0005 | Brassicales | Eudicotyledones | phytozome |
| *Alyssum linifolium* | Alyli.0031s0213 | Brassicales | Eudicotyledones | phytozome |
| *Alyssum linifolium* | Alyli.0216s0006 | Brassicales | Eudicotyledones | phytozome |
| *Amaranthus hypochondriacus* | AH006150 | Caryophyllales | Eudicotyledones | phytozome |
| *Amaranthus hypochondriacus* | AH001650 | Caryophyllales | Eudicotyledones | phytozome |
| *Amborella trichopoda* | evm_27.model.AmTr_v1.0_scaff<br>old00177.28 | Amborellales | Amborellales | phytozome |
| *Amborella trichopoda* | evm_27.model.AmTr_v1.0_scaff<br>old00051.97 | Amborellales | Amborellales | phytozome |
| *Anacardium occidentale* | Anaoc.0018s0955 | Sapindales | Eudicotyledones | phytozome |
| *Anacardium occidentale* | Anaoc.0017s0442 | Sapindales | Eudicotyledones | phytozome |
| *Anacardium occidentale* | Anaoc.1291s0006 | Sapindales | Eudicotyledones | phytozome |
| *Anacardium occidentale* | Anaoc.0003s1165 | Sapindales | Eudicotyledones | phytozome |
| *Ananas comosus* | Aco005843 | Poales | Monocotyledones | phytozome |
| *Ananas comosus* | Aco024739 | Poales | Monocotyledones | phytozome |
| *Aquilegia coerulea* | Aqcoe3G057800 | Ranunculales | Eudicotyledones | phytozome |
| *Aquilegia coerulea* | Aqcoe5G142900 | Ranunculales | Eudicotyledones | phytozome |
| *Aquilegia coerulea* | Aqcoe6G311500 | Ranunculales | Eudicotyledones | phytozome |
| *Arabidopsis halleri* | Araha.12280s0002 | Brassicales | Eudicotyledones | phytozome |
| *Arabidopsis halleri* | Araha.2717s0004 | Brassicales | Eudicotyledones | phytozome |
| *Arabidopsis lyrata* | AL7G50930 | Brassicales | Eudicotyledones | phytozome |
| *Arabidopsis lyrata* | AL3G32830 | Brassicales | Eudicotyledones | phytozome |
| *Arabidopsis lyrata* | AL5G30230 | Brassicales | Eudicotyledones | phytozome |
| *Arabidopsis thaliana* | At5g39400 | Brassicales | Eudicotyledones | phytozome |
| *Arabidopsis thaliana* | At3g19420 | Brassicales | Eudicotyledones | phytozome |
| *Arabidopsis thaliana* | At3g50110 | Brassicales | Eudicotyledones | phytozome |
| *Arachis hypogaea* | arahy.Tifrunner.gnm1.ann1.854<br>N7M | Fabales | Eudicotyledones | phytozome |
| *Arachis hypogaea* | arahy.Tifrunner.gnm1.ann1.PFF<br>9RU | Fabales | Eudicotyledones | phytozome |
| *Arachis hypogaea* | arahy.Tifrunner.gnm1.ann1.132<br>VHE | Fabales | Eudicotyledones | phytozome |
| *Arachis hypogaea* | arahy.Tifrunner.gnm1.ann1.69L<br>NNJ | Fabales | Eudicotyledones | phytozome |
| *Arachis hypogaea* | arahy.Tifrunner.gnm1.ann1.AK4<br>RCU | Fabales | Eudicotyledones | phytozome |
| *Asparagus officinalis* | evm.model.AsparagusV1_08.13<br>99 | Asparagales | Monocotyledones | phytozome |
| *Asparagus officinalis* | evm.model.AsparagusV1_07.85<br>7 | Asparagales | Monocotyledones | phytozome |
| *Asparagus officinalis* | evm.model.AsparagusV1_08.32<br>54 | Asparagales | Monocotyledones | phytozome |
| *Aureococcus anophagefferens* | AURANDRAFT_26423 | Pelagomonadales | Heterokonts | NCBI |
| *Aureococcus anophagefferens* | AURANDRAFT_71576 | Pelagomonadales | Heterokonts | NCBI |
| *Aureococcus anophagefferens* | AURANDRAFT_70790 | Pelagomonadales | Heterokonts | NCBI |
| *Beta vulgaris* | EL10Ac9g22309 | Caryophyllales | Eudicotyledones | phytozome |
| *Beta vulgaris* | EL10Ac6g15699 | Caryophyllales | Eudicotyledones | phytozome |
| *Betula platyphylla* | BPChr11G05686 | Fagales | Eudicotyledones | phytozome |
| *Betula platyphylla* | BPChr06G22535 | Fagales | Eudicotyledones | phytozome |
| *Boechnera stricta* | Bostr.25849s0003 | Brassicales | Eudicotyledones | phytozome |
| *Boechnera stricta* | Bostr.19424s0720 | Brassicales | Eudicotyledones | phytozome |
| *Boechnera stricta* | Bostr.6864s0231 | Brassicales | Eudicotyledones | phytozome |
| *Brachypodium distachyon* | Bradi4g08080 | Poales | Monocotyledones | phytozome |
| *Brachypodium hybridum* | Brahy.D04G0113800 | Poales | Monocotyledones | phytozome |
| *Brachypodium hybridum* | Brahy.S10G0102100 | Poales | Monocotyledones | phytozome |
| *Brachypodium sylvaticum* | Brasy5G104500 | Poales | Monocotyledones | phytozome |
| *Brassica oleracea capitata* | Bol040114 | Brassicales | Eudicotyledones | phytozome |
| *Brassica oleracea capitata* | Bol018098 | Brassicales | Eudicotyledones | phytozome |
| *Brassica oleracea capitata* | Bol031011 | Brassicales | Eudicotyledones | phytozome |
| *Brassica oleracea capitata* | Bol016970 | Brassicales | Eudicotyledones | phytozome |
| *Brassica rapa* | Brara.A02880 | Brassicales | Eudicotyledones | phytozome |
| *Brassica rapa* | Brara.E02262 | Brassicales | Eudicotyledones | phytozome |
| *Brassica rapa* | Brara.C04308 | Brassicales | Eudicotyledones | phytozome |
| *Cakile maritima* | Camar.1507s0005 | Brassicales | Eudicotyledones | phytozome |
| *Cakile maritima* | Camar.0166s0020 | Brassicales | Eudicotyledones | phytozome |
| *Cakile maritima* | Camar.0140s0022 | Brassicales | Eudicotyledones | phytozome |
| *Cakile maritima* | Camar.0046s0044 | Brassicales | Eudicotyledones | phytozome |
| *Capsella grandiflora* | Cagra.12117s0008 | Brassicales | Eudicotyledones | phytozome |
| *Capsella grandiflora* | Cagra.1211s0012 | Brassicales | Eudicotyledones | phytozome |
| *Capsella grandiflora* | Cagra.26626s0001 | Brassicales | Eudicotyledones | phytozome |
| *Capsella rubella* | Carub.0007s3654 | Brassicales | Eudicotyledones | phytozome |
| *Capsella rubella* | Carub.0003s1923 | Brassicales | Eudicotyledones | phytozome |
| *Capsella rubella* | Carub.0005s1645 | Brassicales | Eudicotyledones | phytozome |
| *Carex littledalei* | FCM35_KLT03997 | Cyperales | Monocotyledones | NCBI |
| *Carex littledalei* | FCM35_KLT07398 | Cyperales | Monocotyledones | NCBI |
| *Carica papaya* | evm.model.supercontig_14.140 | Brassicales | Eudicotyledones | phytozome |
| *Carica papaya* | evm.model.supercontig_200.15 | Brassicales | Eudicotyledones | phytozome |
| *Carya illinoensis* | Caril.03G085200 | Juglandales |  | phytozome |
| *Carya illinoensis* | Caril.04G059000 | Juglandales |  | phytozome |
| *Carya illinoensis* | Caril.01G012500 | Juglandales |  | phytozome |
| *Carya illinoensis* | Caril.06G171800 | Juglandales |  | phytozome |
| *Carya illinoensis* | Caril.05G009200 | Juglandales |  | phytozome |
| *Castanea dentata* | Caden.01G186200 | Fagales | Eudicotyledones | phytozome |
| *Castanea dentata* | Caden.02G186200 | Fagales | Eudicotyledones | phytozome |
| *Castanea dentata* | Caden.07G119900 | Fagales | Eudicotyledones | phytozome |
| *Caulanthus amplexicaulis* | Caamp.0044s0232 | Brassicales | Eudicotyledones | phytozome |
| *Caulanthus amplexicaulis* | Caamp.1041s0897 | Brassicales | Eudicotyledones | phytozome |
| *Caulanthus amplexicaulis* | Caamp.1039s1127 | Brassicales | Eudicotyledones | phytozome |
| *Caulanthus amplexicaulis* | Caamp.0051s0388 | Brassicales | Eudicotyledones | phytozome |
| *Caulanthus amplexicaulis* | Caamp.0078s0093 | Brassicales | Eudicotyledones | phytozome |
| *Caulanthus amplexicaulis* | Caamp.0026s0499 | Brassicales | Eudicotyledones | phytozome |
| *Ceratodon purpureus* | CepurGG1.8G115300 | Dicranales | Bryophytes | phytozome |
| *Ceratodon purpureus* | CepurGG1.11G093000 | Dicranales | Bryophytes | phytozome |
| *Ceratopteris richardii* | Ceric.25G072600 | Polypodiales | Monilophytes | phytozome |
| *Ceratopteris richardii* | Ceric.16G021900 | Polypodiales | Monilophytes | phytozome |
| *Chara braunii* | CBR_g29865 | Charales | Charophytes | NCBI |
| *Chara braunii* | CBR_g30221 | Charales | Charophytes | NCBI |
| *Chenopodium quinoa* | AUR62013248 | Caryophyllales | Eudicotyledones | phytozome |
| *Chenopodium quinoa* | AUR62010267 | Caryophyllales | Eudicotyledones | phytozome |
| *Chlamydomonas reinhardtii* | Cre06.g308400 | Chlamydomonales | Chlorophytes | phytozome |
| *Chromochloris zofingensis* | Cz03g22130.t1 | Sphaeropleales | Chlorophytes | phytozome |
| *Chrysochromulina tobinii* | Ctob_005637 | Prymniales | Haptophytes | NCBI |
| *Chrysochromulina tobinii* | Ctob_015421 | Prymniales | Haptophytes | NCBI |
| *Chrysochromulina tobinii* | Ctob_003434 | Prymniales | Haptophytes | NCBI |
| *Chrysochromulina tobinii* | Ctob_004562 | Prymniales | Haptophytes | NCBI |
| *Chrysochromulina tobinii* | Ctob_005565 | Prymniales | Haptophytes | NCBI |
| *Cicer arietinum* | Ca_02178 | Fabales | Eudicotyledones | phytozome |
| *Cicer arietinum* | Ca_05590 | Fabales | Eudicotyledones | phytozome |
| *Cinnamomum kanehirae* | CKAN_02766500 | Laurales | Magnoliids | phytozome |
| *Cinnamomum kanehirae* | CKAN_02500000 | Laurales | Magnoliids | phytozome |
| *Cinnamomum kanehirae* | CKAN_01173600 | Laurales | Magnoliids | phytozome |
| *Cinnamomum kanehirae* | CKAN_00608500 | Laurales | Magnoliids | phytozome |
| *Citrus clementina* | Ciclev10011769m | Sapindales | Eudicotyledones | phytozome |
| *Citrus clementina* | Ciclev10025539m | Sapindales | Eudicotyledones | phytozome |
| *Citrus clementina* | Ciclev10028042m | Sapindales | Eudicotyledones | phytozome |
| *Citrus sinensis* | orange1.1g012952m | Sapindales | Eudicotyledones | phytozome |
| *Citrus sinensis* | orange1.1g014325m | Sapindales | Eudicotyledones | phytozome |
| *Citrus sinensis* | orange1.1g037030m | Sapindales | Eudicotyledones | phytozome |
| *Cleome violacea* | Clevi.0004s2083 | Brassicales | Eudicotyledones | phytozome |
| *Cleome violacea* | Clevi.0032s0574 | Brassicales | Eudicotyledones | phytozome |
| *Cleome violacea* | Clevi.0005s2503 | Brassicales | Eudicotyledones | phytozome |
| *Coccomyxa subellipsoidea* | 66502 |  | Chlorophytes | phytozome |
| *Coffea arabica* | evm.model.Scaffold_952.463 | Gentianales | Eudicotyledones | phytozome |
| *Coffea arabica* | evm.model.Scaffold_612.518 | Gentianales | Eudicotyledones | phytozome |
| *Coffea arabica* | evm.model.Scaffold_634.624 | Gentianales | Eudicotyledones | phytozome |
| *Coffea arabica* | evm.model.Scaffold_952.172 | Gentianales | Eudicotyledones | phytozome |
| *Corymbia citriodora* | Cocit.A0156 | Myrtales | Eudicotyledones | phytozome |
| *Corymbia citriodora* | Cocit.L5056 | Myrtales | Eudicotyledones | phytozome |
| *Corymbia citriodora* | Cocit.G0778 | Myrtales | Eudicotyledones | phytozome |
| *Crambe hispanica* | Crahi.1829s0005 | Brassicales | Eudicotyledones | phytozome |
| *Crambe hispanica* | Crahi.0455s0005 | Brassicales | Eudicotyledones | phytozome |
| *Crambe hispanica* | Crahi.0412s0031 | Brassicales | Eudicotyledones | phytozome |
| *Crambe hispanica* | Crahi.0943s0003 | Brassicales | Eudicotyledones | phytozome |
| *Crambe hispanica* | Crahi.0276s0014 | Brassicales | Eudicotyledones | phytozome |
| *Cucumis sativus* | Cucsa.338540 | Cucurbitales | Eudicotyledones | phytozome |
| *Daucus carota* | DCAR_001632 | Apiales | Eudicotyledones | phytozome |
| *Daucus carota* | DCAR_029077 | Apiales | Eudicotyledones | phytozome |
| *Daucus carota* | DCAR_010857 | Apiales | Eudicotyledones | phytozome |
| *Descurainia sophioides* | Desop.0231s0613 | Brassicales | Eudicotyledones | phytozome |
| *Descurainia sophioides* | Desop.0248s0666 | Brassicales | Eudicotyledones | phytozome |
| *Descurainia sophioides* | Desop.0207s0242 | Brassicales | Eudicotyledones | phytozome |
| *Dichanthelium oligosanthes* | BAE44_0008647 | Poales | Monocotyledones | NCBI |
| *Dioscorea alata* | Dioal.19G152700 | Dioscoreales | Monocotyledones | phytozome |
| *Diptychocarpus strictus* | Distr.0012s22625 | Brassicales | Eudicotyledones | phytozome |
| *Diptychocarpus strictus* | Distr.0005s223300 | Brassicales | Eudicotyledones | phytozome |
| *Diptychocarpus strictus* | Distr.0006s170500 | Brassicales | Eudicotyledones | phytozome |
| *Ectocarpus siliculosus* | Esi_0301_0020 | Ectocarpales | Heterokonts | NCBI |
| *Ectocarpus siliculosus* | Esi_0147_0077 | Ectocarpales | Heterokonts | NCBI |
| *Ectocarpus siliculosus* | Esi_0147_0079 | Ectocarpales | Heterokonts | NCBI |
| *Elaeis guineensis* | LOC105056003 | Arecales | Monocotyledones | NCBI |
| *Elaeis guineensis* | LOC105045483 | Arecales | Monocotyledones | NCBI |
| *Elaeis guineensis* | LOC105035125 | Arecales | Monocotyledones | NCBI |
| *Eleusine coracana* | ELECO.r07.5BG0436130 | Poales | Monocotyledones | phytozome |
| *Eleusine coracana* | ELECO.r07.5AG0388310 | Poales | Monocotyledones | phytozome |
| *Emiliania huxleyi* | EMIHUDDRAFT_195717 | Isochrysidales | Haptophytes | NCBI |
| *Emiliania huxleyi* | EMIHUDDRAFT_437207 | Isochrysidales | Haptophytes | NCBI |
| *Eragrostis curvula* | EJB05_06262 | Cyperales | Monocotyledones | NCBI |
| *Eragrostis curvula* | EJB05_01954 | Cyperales | Monocotyledones | NCBI |
| *Eruca vesicaria* | Eruve.3288s0001 | Brassicales | Eudicotyledones | phytozome |
| *Eruca vesicaria* | Eruve.2621s0007 | Brassicales | Eudicotyledones | phytozome |
| *Eruca vesicaria* | Eruve.2403s0005 | Brassicales | Eudicotyledones | phytozome |
| *Eruca vesicaria* | Eruve.0429s0037 | Brassicales | Eudicotyledones | phytozome |
| *Eruca vesicaria* | Eruve.1019s0020 | Brassicales | Eudicotyledones | phytozome |
| *Eruca vesicaria* | Eruve.1393s0007 | Brassicales | Eudicotyledones | phytozome |
| *Eucalyptus grandis* | Eucgr.J01367 | Myrtales | Eudicotyledones | phytozome |
| *Eucalyptus grandis* | Eucgr.G01374 | Myrtales | Eudicotyledones | phytozome |
| *Eucalyptus grandis* | Eucgr.I00804 | Myrtales | Eudicotyledones | phytozome |
| *Euclidium syriacum* | Eusyr.0002s0385 | Brassicales | Eudicotyledones | phytozome |
| *Euclidium syriacum* | Eusyr.0017s0515 | Brassicales | Eudicotyledones | phytozome |
| *Euclidium syriacum* | Eusyr.0120s0222 | Brassicales | Eudicotyledones | phytozome |
| *Eutrema salsugineum* | Thhalv10027783m | Brassicales | Eudicotyledones | phytozome |
| *Eutrema salsugineum* | Thhalv10020332m | Brassicales | Eudicotyledones | phytozome |
| *Eutrema salsugineum* | Thhalv10010193m | Brassicales | Eudicotyledones | phytozome |
| *Fragaria vesca* | FvH4_6g47530 | Rosales | Eudicotyledones | phytozome |
| *Fragaria vesca* | FvH4_2g05490 | Rosales | Eudicotyledones | phytozome |
| *Fragaria vesca* | FvH4_1g16580 | Rosales | Eudicotyledones | phytozome |
| *Ginkgo biloba* | GBI00005494 | Ginkgoales | Acrogymnosperms | Gymno<br>plaza 1.0 |
| *Glycine max* | GlysoPI483463.08G218300 | Fabales | Eudicotyledones | phytozome |
| *Glycine max* | GlysoPI483463.01G151800 | Fabales | Eudicotyledones | phytozome |
| *Glycine max* | GlysoPI483463.11G048000 | Fabales | Eudicotyledones | phytozome |
| *Glycine max* | GlysoPI483463.10G224200 | Fabales | Eudicotyledones | phytozome |
| *Glycine max* | GlysoPI483463.20G094900 | Fabales | Eudicotyledones | phytozome |
| *Gnetum montanum* | GMO00031901 | Gnetales | Acrogymnosperms | Gymno<br>plaza 1.0 |
| *Gossypium bardadense* | Gobar.A11G078400 | Malvales | Eudicotyledones | phytozome |
| *Gossypium bardadense* | Gobar.D11G079100 | Malvales | Eudicotyledones | phytozome |
| *Gossypium bardadense* | Gobar.D11G053600 | Malvales | Eudicotyledones | phytozome |
| *Gossypium bardadense* | Gobar.A11G053700 | Malvales | Eudicotyledones | phytozome |
| *Gossypium bardadense* | Gobar.D02G241200 | Malvales | Eudicotyledones | phytozome |
| *Gossypium bardadense* | Gobar.A03G202000 | Malvales | Eudicotyledones | phytozome |
| *Gossypium bardadense* | Gobar.A12G022900 | Malvales | Eudicotyledones | phytozome |
| *Gossypium bardadense* | Gobar.D12G024700 | Malvales | Eudicotyledones | phytozome |
| *Gossypium bardadense* | Gobar.D01G007700 | Malvales | Eudicotyledones | phytozome |
| *Gossypium bardadense* | Gobar.A01G007300 | Malvales | Eudicotyledones | phytozome |
| *Gossypium hirsutum* | Gohir.A11G071400 | Malvales | Eudicotyledones | phytozome |
| *Gossypium hirsutum* | Gohir.D11G050950 | Malvales | Eudicotyledones | phytozome |
| *Gossypium hirsutum* | Gohir.A11G047500 | Malvales | Eudicotyledones | phytozome |
| *Gossypium hirsutum* | Gohir.D11G051100 | Malvales | Eudicotyledones | phytozome |
| *Guillardia theta* | GUITHDRAFT_146552 | Pyrenomonadales | Cryptomonades | NCBI |
| *Guillardia theta* | GUITHDRAFT_158972 | Pyrenomonadales | Cryptomonades | NCBI |
| *Guillardia theta* | GUITHDRAFT_136489 | Pyrenomonadales | Cryptomonades | NCBI |
| *Helianthus annuus* | HanXRQChr15g0473081 | Asterales | Eudicotyledones | phytozome |
| *Helianthus annuus* | HanXRQChr03g0070181 | Asterales | Eudicotyledones | phytozome |
| *Helianthus annuus* | HanXRQChr04g0128111 | Asterales | Eudicotyledones | phytozome |
| *Helianthus annuus* | HanXRQChr11g0348401 | Asterales | Eudicotyledones | phytozome |
| *Hydrangea quercifolia* | Hyque.05G148600 | Rosales | Eudicotyledones | phytozome |
| *Hydrangea quercifolia* | Hyque.03G170500 | Rosales | Eudicotyledones | phytozome |
| *Hydrangea quercifolia* | Hyque.13G030400 | Rosales | Eudicotyledones | phytozome |
| *Iberis amara* | Ibeam.3529s0003 | Brassicales | Eudicotyledones | phytozome |
| *Iberis amara* | Ibeam.1232s0005 | Brassicales | Eudicotyledones | phytozome |
| *Iberis amara* | Ibeam.6756s0002 | Brassicales | Eudicotyledones | phytozome |
| *Iberis amara* | Ibeam.3246s0002 | Brassicales | Eudicotyledones | phytozome |
| *Isatis tinctoria* | Isati.8565s0004 | Brassicales | Eudicotyledones | phytozome |
| *Isatis tinctoria* | Isati.0832s0027 | Brassicales | Eudicotyledones | phytozome |
| *Isatis tinctoria* | Isati.0178s0011 | Brassicales | Eudicotyledones | phytozome |
| *Isatis tinctoria* | Isati.1336s0016 | Brassicales | Eudicotyledones | phytozome |
| *Isatis tinctoria* | Isati.1514s0004 | Brassicales | Eudicotyledones | phytozome |
| *Joinvillea ascendens* | Joasc.14G102000 | Poales | Monocotyledones | phytozome |
| *Kalanchoe fedtschenkoi* | Kaladp0026s0043 | Saxifragales | Eudicotyledones | phytozome |
| *Kalanchoe fedtschenkoi* | Kaladp0015s0211 | Saxifragales | Eudicotyledones | phytozome |
| *Kalanchoe fedtschenkoi* | Kaladp0058s0422 | Saxifragales | Eudicotyledones | phytozome |
| *Kalanchoe fedtschenkoi* | Kaladp0028s0094 | Saxifragales | Eudicotyledones | phytozome |
| *Klebsormidium nitens* | GAQ85806 | Klebsormidiales | Charophytes | NCBI |
| *Klebsormidium nitens* | GAQ91864 | Klebsormidiales | Charophytes | NCBI |
| *Klebsormidium nitens* | GAQ91212 | Klebsormidiales | Charophytes | NCBI |
| *Klebsormidium nitens* | GAQ79654 | Klebsormidiales | Charophytes | NCBI |
| *Lactuca sativa* | Lsat_1_v5_gn_3_98660 | Asterales | Eudicotyledones | phytozome |
| *Lactuca sativa* | Lsat_1_v5_gn_3_98880 | Asterales | Eudicotyledones | phytozome |
| *Lactuca sativa* | Lsat_1_v5_gn_7_89141 | Asterales | Eudicotyledones | phytozome |
| *Lactuca sativa* | Lsat_1_v5_gn_3_98761 | Asterales | Eudicotyledones | phytozome |
| *Lactuca sativa* | Lsat_1_v5_gn_3_98721 | Asterales | Eudicotyledones | phytozome |
| *Lactuca sativa* | Lsat_1_v5_gn_3_90360 | Asterales | Eudicotyledones | phytozome |
| *Lactuca sativa* | Lsat_1_v5_gn_9_111380 | Asterales | Eudicotyledones | phytozome |
| *Lactuca sativa* | Lsat_1_v5_gn_5_70961 | Asterales | Eudicotyledones | phytozome |
| *Lepidium sativum* | Lesat.0070s0837 | Brassicales | Eudicotyledones | phytozome |
| *Lepidium sativum* | Lesat.0041s0230 | Brassicales | Eudicotyledones | phytozome |
| *Lepidium sativum* | Lesat.0086s0034 | Brassicales | Eudicotyledones | phytozome |
| *Lepidium sativum* | Lesat.0019s0255 | Brassicales | Eudicotyledones | phytozome |
| *Lepidium sativum* | Lesat.0013s0306 | Brassicales | Eudicotyledones | phytozome |
| *Lepidium sativum* | Lesat.0024s0016 | Brassicales | Eudicotyledones | phytozome |
| *Lindenbergia philippensis* | Liphi.09G031000 | Lamiales | Eudicotyledones | phytozome |
| *Lindenbergia philippensis* | Liphi.02G151000 | Lamiales | Eudicotyledones | phytozome |
| *Lindenbergia philippensis* | Liphi.09G060300 | Lamiales | Eudicotyledones | phytozome |
| *Linum usitatissimum* | Lus10027876 | Malpighiales | Eudicotyledones | phytozome |
| *Linum usitatissimum* | Lus10002826 | Malpighiales | Eudicotyledones | phytozome |
| *Lotus japonicus* | Lj1g0022114 | Fabales | Eudicotyledones | phytozome |
| *Lotus japonicus* | Lj2g0026106 | Fabales | Eudicotyledones | phytozome |
| *Lotus japonicus* | Lj5g0017669 | Fabales | Eudicotyledones | phytozome |
| *Lunaria annua* | Luann.0281s0031 | Brassicales | Eudicotyledones | phytozome |
| *Lunaria annua* | Luann.0007s0091 | Brassicales | Eudicotyledones | phytozome |
| *Lunaria annua* | Luann.0026s0012 | Brassicales | Eudicotyledones | phytozome |
| *Lunaria annua* | Luann.0006s0072 | Brassicales | Eudicotyledones | phytozome |
| *Lupinus albus* | Lalb_Ch17g0337091 | Fabales | Eudicotyledones | phytozome |
| *Lupinus albus* | Lalb_Ch04g0258551 | Fabales | Eudicotyledones | phytozome |
| *Lupinus albus* | Lalb_Ch16g0378681 | Fabales | Eudicotyledones | phytozome |
| *Lupinus albus* | Lalb_Ch21g0308431 | Fabales | Eudicotyledones | phytozome |
| *Malcomia maritima* | Mamar.0082s0142 | Brassicales | Eudicotyledones | phytozome |
| *Malcomia maritima* | Mamar.0029s0743 | Brassicales | Eudicotyledones | phytozome |
| *Malcomia maritima* | Mamar.0003s0492 | Brassicales | Eudicotyledones | phytozome |
| *Malus domestica* | MD17G1056300 | Rosales | Eudicotyledones | phytozome |
| *Malus domestica* | MD09G1060900 | Rosales | Eudicotyledones | phytozome |
| *Malus domestica* | MD02G1177900 | Rosales | Eudicotyledones | phytozome |
| *Malus domestica* | MD15G1287800 | Rosales | Eudicotyledones | phytozome |
| *Malus domestica* | MD05G1087100 | Rosales | Eudicotyledones | phytozome |
| *Manihot esculenta* | Manes.02G050801 | Malpighiales | Eudicotyledones | phytozome |
| *Manihot esculenta* | Manes.12G108000 | Malpighiales | Eudicotyledones | phytozome |
| *Manihot esculenta* | Manes.10G077300 | Malpighiales | Eudicotyledones | phytozome |
| *Marchantia polymorpha* | Mapoly0058s0093 | Marchantiales | Marchantiophytes | phytozome |
| *Marchantia polymorpha* | Mapoly0016s0179 | Marchantiales | Marchantiophytes | phytozome |
| *Marchantia polymorpha* | Mapoly0117s0012 | Marchantiales | Marchantiophytes | phytozome |
| *Medicago truncatula* | Medtr7g012250 | Fabales | Eudicotyledones | phytozome |
| *Medicago truncatula* | Medtr5g016550 | Fabales | Eudicotyledones | phytozome |
| *Medicago truncatula* | Medtr1g107540 | Fabales | Eudicotyledones | phytozome |
| *Mimulus guttatus* | Migut.H00656 | Lamiales | Eudicotyledones | phytozome |
| *Mimulus guttatus* | Migut.H00402 | Lamiales | Eudicotyledones | phytozome |
| *Mimulus guttatus* | Migut.B00375 | Lamiales | Eudicotyledones | phytozome |
| *Miscanthus sinensis* | Misin03G029900 | Cyperales | Monocotyledones | phytozome |
| *Miscanthus sinensis* | Misin04G012900 | Cyperales | Monocotyledones | phytozome |
| *Musa acuminata* | GSMUA_Achr10P29220_001 | Zingiberales | Monocotyledones | phytozome |
| *Musa acuminata* | GSMUA_Achr7P21640_001 | Zingiberales | Monocotyledones | phytozome |
| *Musa acuminata* | GSMUA_Achr5P07740_001 | Zingiberales | Monocotyledones | phytozome |
| *Musa balbisiana* | C4D60_Mb05t09990 | Zingiberales | Monocotyledones | NCBI |
| *Musa balbisiana* | C4D60_Mb07t05750 | Zingiberales | Monocotyledones | NCBI |
| *Musa balbisiana* | C4D60_Mb10t02290 | Zingiberales | Monocotyledones | NCBI |
| *Myagrurn perfoliatum* | Myper.0019s0558 | Brassicales | Eudicotyledones | phytozome |
| *Myagrurn perfoliatum* | Myper.0009s1500 | Brassicales | Eudicotyledones | phytozome |
| *Myagrurn perfoliatum* | Myper.0005s1433 | Brassicales | Eudicotyledones | phytozome |
| *Nelumbo nucifera* | LOC104598017 | Proteales | Eudicotyledones | NCBI |
| *Nelumbo nucifera* | LOC104589843 | Proteales | Eudicotyledones | NCBI |
| *Nelumbo nucifera* | LOC104605416 | Proteales | Eudicotyledones | NCBI |
| *Nymphaea colorata* | Nycol.I01022 | Nymphaeales | Nymphaeales | phytozome |
| *Nymphaea colorata* | Nycol.C00833 | Nymphaeales | Nymphaeales | phytozome |
| *Olea europaea* | Oeu062142 | Lamiales | Eudicotyledones | phytozome |
| *Olea europaea* | Oeu014788 | Lamiales | Eudicotyledones | phytozome |
| *Olea europaea* | Oeu001763 | Lamiales | Eudicotyledones | phytozome |
| *Olea europaea* | Oeu044292 | Lamiales | Eudicotyledones | phytozome |
| *Oryza sativa* | LOC_Os12g21890 | Poales | Monocotyledones | phytozome |
| *Panicum hallii* | Pahal.2G380700 | Poales | Monocotyledones | phytozome |
| *Panicum virgatum* | Pavir.2KG446900 | Poales | Monocotyledones | phytozome |
| *Panicum virgatum* | Pavir.2NG500600 | Poales | Monocotyledones | phytozome |
| *Paspalum vaginatum* | Pavag02G285600 | Poales | Monocotyledones | phytozome |
| *Pharus latifolius* | Phala.10G061700 | Poales | Monocotyledones | phytozome |
| *Phaseolus vulgaris* | Phvul.008G036300 | Fabales | Eudicotyledones | phytozome |
| *Phaseolus vulgaris* | Phvul.002G015700 | Fabales | Eudicotyledones | phytozome |
| *Phaseolus vulgaris* | Phvul.007G038300 | Fabales | Eudicotyledones | phytozome |
| *Phoenix dactylifera* | LOC103711183 | Arecales | Monocotyledones | NCBI |
| *Phoenix dactylifera* | LOC103706227 | Arecales | Monocotyledones | NCBI |
| *Phoenix dactylifera* | LOC103705030 | Arecales | Monocotyledones | NCBI |
| *Physcomitrella patens* | Pp3c19_18320V3 | Funariales | Bryophytes | phytozome |
| *Physcomitrella patens* | Pp3c22_10340V3 | Funariales | Bryophytes | phytozome |
| *Physcomitrella patens* | Pp3c21_8410V3 | Funariales | Bryophytes | phytozome |
| *Physcomitrella patens* | Pp3c22_14420V3 | Funariales | Bryophytes | phytozome |
| *Picea glauca* | PGL00015278 | Pinales | Acrogymnosperms | Gymno<br>plaza 1.0 |
| *Pinus pinaster* | PPI00036745 | Pinales | Acrogymnosperms | Gymno<br>plaza 1.0 |
| *Pinus pinaster* | PPI00005815 | Pinales | Acrogymnosperms | Gymno<br>plaza 1.0 |
| *Pinus pinaster* | PPI00034805 | Pinales | Acrogymnosperms | Gymno<br>plaza 1.0 |
| *Pinus sylvestris* | PSY00021791 | Pinales | Acrogymnosperms | Gymno<br>plaza 1.0 |
| *Pinus sylvestris* | PSY00016716 | Pinales | Acrogymnosperms | Gymno<br>plaza 1.0 |
| *Pinus taeda* | PTA00064026 | Pinales | Acrogymnosperms | Gymno<br>plaza 1.0 |
| *Pinus taeda* | PTA00032593 | Pinales | Acrogymnosperms | Gymno<br>plaza 1.0 |
| *Pinus taeda* | PTA00009584 | Pinales | Acrogymnosperms | Gymno<br>plaza 1.0 |
| *Poncirus trifoliata* | Ptrif.0006s2220 | Sapindales | Eudicotyledones | phytozome |
| *Poncirus trifoliata* | Ptrif.0007s0475 | Sapindales | Eudicotyledones | phytozome |
| *Poncirus trifoliata* | Ptrif.0008s0442 | Sapindales | Eudicotyledones | phytozome |
| *Populus deltoides* | Podel.17G093700 | Malpighiales | Eudicotyledones | phytozome |
| *Populus deltoides* | Podel.07G056300 | Malpighiales | Eudicotyledones | phytozome |
| *Populus deltoides* | Podel.01G321200 | Malpighiales | Eudicotyledones | phytozome |
| *Populus deltoides* | Podel.09G100700 | Malpighiales | Eudicotyledones | phytozome |
| *Populus trichocarpa* | Potri.017G089700 | Malpighiales | Eudicotyledones | phytozome |
| *Populus trichocarpa* | Potri.007G047900 | Malpighiales | Eudicotyledones | phytozome |
| *Populus trichocarpa* | Potri.001G302000 | Malpighiales | Eudicotyledones | phytozome |
| *Populus trichocarpa* | Potri.009G098000 | Malpighiales | Eudicotyledones | phytozome |
| *Portulaca amilis* | FUN_051737 | Caryophyllales | Eudicotyledones | phytozome |
| *Portulaca amilis* | FUN_052803 | Caryophyllales | Eudicotyledones | phytozome |
| *Portulaca amilis* | FUN_047201 | Caryophyllales | Eudicotyledones | phytozome |
| *Prunus persica* | Prupe.3G259200 | Rosales | Eudicotyledones | phytozome |
| *Prunus persica* | Prupe.6G230100 | Rosales | Eudicotyledones | phytozome |
| *Prunus persica* | Prupe.8G131700 | Rosales | Eudicotyledones | phytozome |
| *Pseudotsuga menziesii* | PME00026064 | Pinales | Acrogymnosperms | Gymno<br>plaza 1.0 |
| *Pseudotsuga menziesii* | PME00001365 | Pinales | Acrogymnosperms | Gymno<br>plaza 1.0 |
| *Pseudotsuga menziesii* | PME00077364 | Pinales | Acrogymnosperms | Gymno<br>plaza 1.0 |
| *Quercus rubra* | Qurub.02G186400 | Fagales | Eudicotyledones | phytozome |
| *Quercus rubra* | Qurub.11G098700 | Fagales | Eudicotyledones | phytozome |
| *Quercus rubra* | Qurub.06G075200 | Fagales | Eudicotyledones | phytozome |
| *Ricinus communis* | 29970.m000976 | Malpighiales | Eudicotyledones | phytozome |
| *Ricinus communis* | 29801.m003123 | Malpighiales | Eudicotyledones | phytozome |
| *Rorippa islandica* | Roisl.0070s0018 | Brassicales | Eudicotyledones | phytozome |
| *Rorippa islandica* | Roisl.0046s0950 | Brassicales | Eudicotyledones | phytozome |
| *Salix purpurea* | Sapur.017G073600 | Salicales | Eudicotyledones | phytozome |
| *Salix purpurea* | Sapur.007G045200 | Salicales | Eudicotyledones | phytozome |
| *Salix purpurea* | Sapur.016G181600 | Salicales | Eudicotyledones | phytozome |
| *Salix purpurea* | Sapur.009G076600 | Salicales | Eudicotyledones | phytozome |
| *Schrenkiella parvula* | Sp7g02580 | Brassicales | Eudicotyledones | phytozome |
| *Schrenkiella parvula* | Sp3g17480 | Brassicales | Eudicotyledones | phytozome |
| *Schrenkiella parvula* | Sp5g12010 | Brassicales | Eudicotyledones | phytozome |
| *Selaginella moellendorffii* | 91219 | Selaginellales | Lycophytes | phytozome |
| *Selaginella moellendorffii* | 165134 | Selaginellales | Lycophytes | phytozome |
| *Setaria italica* | Seita.2G322700 | Cyperales | Monocotyledones | phytozome |
| *Setaria viridis* | Sevir.2G334300 | Cyperales | Monocotyledones | phytozome |
| *Sinapis alba* | Sialb.0606s0023 | Brassicales | Eudicotyledones | phytozome |
| *Sinapis alba* | Sialb.0540s0029 | Brassicales | Eudicotyledones | phytozome |
| *Sinapis alba* | Sialb.0766s0022 | Brassicales | Eudicotyledones | phytozome |
| *Sinapis alba* | Sialb.0054s0274 | Brassicales | Eudicotyledones | phytozome |
| *Sinapis alba* | Sialb.0001s0595 | Brassicales | Eudicotyledones | phytozome |
| *Sinapis alba* | Sialb.0672s0053 | Brassicales | Eudicotyledones | phytozome |
| *Solanum lycopersicum* | Solyc03g013310 | Solanales | Eudicotyledones | phytozome |
| *Solanum lycopersicum* | Solyc01g107750 | Solanales | Eudicotyledones | phytozome |
| *Solanum lycopersicum* | Solyc02g093000 | Solanales | Eudicotyledones | phytozome |
| *Solanum tuberosum* | Soltu.DM.02G028260 | Solanales | Eudicotyledones | phytozome |
| *Solanum tuberosum* | Soltu.DM.03G007160 | Solanales | Eudicotyledones | phytozome |
| *Solanum tuberosum* | Soltu.DM.01G047110 | Solanales | Eudicotyledones | phytozome |
| *Sorghum bicolor* | Sobic.002G008800 | Poales | Monocotyledones | phytozome |
| *Sphagnum fallax* | Sphfalx16G057000 | Sphagnales | Bryophytes | phytozome |
| *Sphagnum fallax* | Sphfalx17G006600 | Sphagnales | Bryophytes | phytozome |
| *Sphagnum fallax* | Sphfalx18G079700 | Sphagnales | Bryophytes | phytozome |
| *Sphagnum fallax* | Sphfalx13G023500 | Sphagnales | Bryophytes | phytozome |
| *Sphagnum fallax* | Sphfalx14G024100 | Sphagnales | Bryophytes | phytozome |
| *Sphagnum fallax* | Sphfalx06G107500 | Sphagnales | Bryophytes | phytozome |
| *Sphagnum magellanicum* | Sphmag16G056400 | Sphagnales | Bryophytes | phytozome |
| *Sphagnum magellanicum* | Sphmag17G006400 | Sphagnales | Bryophytes | phytozome |
| *Sphagnum magellanicum* | Sphmag06G111300 | Sphagnales | Bryophytes | phytozome |
| *Sphagnum magellanicum* | Sphmag18G019000 | Sphagnales | Bryophytes | phytozome |
| *Sphagnum magellanicum* | Sphmag13G020000 | Sphagnales | Bryophytes | phytozome |
| *Sphagnum magellanicum* | Sphmag14G023700 | Sphagnales | Bryophytes | phytozome |
| *Spinacia oleracea* | Spov3_C0007.00101 | Caryophyllales | Eudicotyledones | phytozome |
| *Spinacia oleracea* | Spov3_chr1.02493 | Caryophyllales | Eudicotyledones | phytozome |
| *Spirodela polyrhiza* | Spipo2G0093300 | Alismatales | Monocotyledones | phytozome |
| *Spirodela polyrhiza* | Spipo4G0005500 | Alismatales | Monocotyledones | phytozome |
| *Stanleya pinnata* | Stapi.1453s0002 | Brassicales | Eudicotyledones | phytozome |
| *Stanleya pinnata* | Stapi.1874s0008 | Brassicales | Eudicotyledones | phytozome |
| *Stanleya pinnata* | Stapi.1161s0006 | Brassicales | Eudicotyledones | phytozome |
| *Stanleya pinnata* | Stapi.0737s0004 | Brassicales | Eudicotyledones | phytozome |
| *Stanleya pinnata* | Stapi.0692s0011 | Brassicales | Eudicotyledones | phytozome |
| *Taxus baccata* | TBA00001628 | Taxales | Acrogymnosperms | Gymno plaza 1.0 |
| *Taxus baccata* | TBA00011682 | Taxales | Acrogymnosperms | Gymno plaza 1.0 |
| *Theobroma cacao* | Thecc.04G090400 | Malvales | Eudicotyledones | phytozome |
| *Theobroma cacao* | Thecc.01G062500 | Malvales | Eudicotyledones | phytozome |
| *Theobroma cacao* | Thecc.02G105700 | Malvales | Eudicotyledones | phytozome |
| *Thinopyrum intermedium* | Thint.15G0279400 | Poales | Monocotyledones | phytozome |
| *Thinopyrum intermedium* | Thint.13G0199200 | Poales | Monocotyledones | phytozome |
| *Thinopyrum intermedium* | Thint.14G0237600 | Poales | Monocotyledones | phytozome |
| *Thlaspi arvense* | Thlar.0083s0010 | Brassicales | Eudicotyledones | phytozome |
| *Thlaspi arvense* | Thlar.0014s0526 | Brassicales | Eudicotyledones | phytozome |
| *Thlaspi arvense* | Thlar.0021s1378 | Brassicales | Eudicotyledones | phytozome |
| *Thuja plicata* | Thupl.29382416s0045 | Pinales | Acrogymnosperms | phytozome |
| *Thuja plicata* | Thupl.29377609s0023 | Pinales | Acrogymnosperms | phytozome |
| *Trifolium pratense* | Tp57577_TGAC_v2_mRNA34577 | Fabales | Eudicotyledones | phytozome |
| *Trifolium pratense* | Tp57577_TGAC_v2_mRNA27810 | Fabales | Eudicotyledones | phytozome |
| *Trifolium pratense* | Tp57577_TGAC_v2_mRNA14329 | Fabales | Eudicotyledones | phytozome |
| *Urochloa fusca* | Urofu.1G032300 | Poales | Monocotyledones | phytozome |
| *Urochloa fusca* | Urofu.2G334900 | Poales | Monocotyledones | phytozome |
| *Vigna unguiculata* | Vigun05g037300 | Fabales | Eudicotyledones | phytozome |
| *Vigna unguiculata* | Vigun07g259200 | Fabales | Eudicotyledones | phytozome |
| *Vigna unguiculata* | Vigun02g146800 | Fabales | Eudicotyledones | phytozome |
| *Vitis vinifera* | VIT_214s0108g01410 | Vitales | Eudicotyledones | phytozome |
| *Vitis vinifera* | VIT_203s0180g00020 | Vitales | Eudicotyledones | phytozome |
| *Vitis vinifera* | VIT_207s0031g00020 | Vitales | Eudicotyledones | phytozome |
| *Volvox carteri* | Vocar.0030s0065 | Chlamydomonadales | Chlorophytes | phytozome |
| *Zea mays* | Zm00001d007942 | Cyperales | Monocotyledones | phytozome |
| *Zostera marina* | Zosma06g28190 | Alismatales | Monocotyledones | phytozome |
| *Zostera marina* | Zosma01g09840 | Alismatales | Monocotyledones | phytozome |

**Supplementary Table 2:**
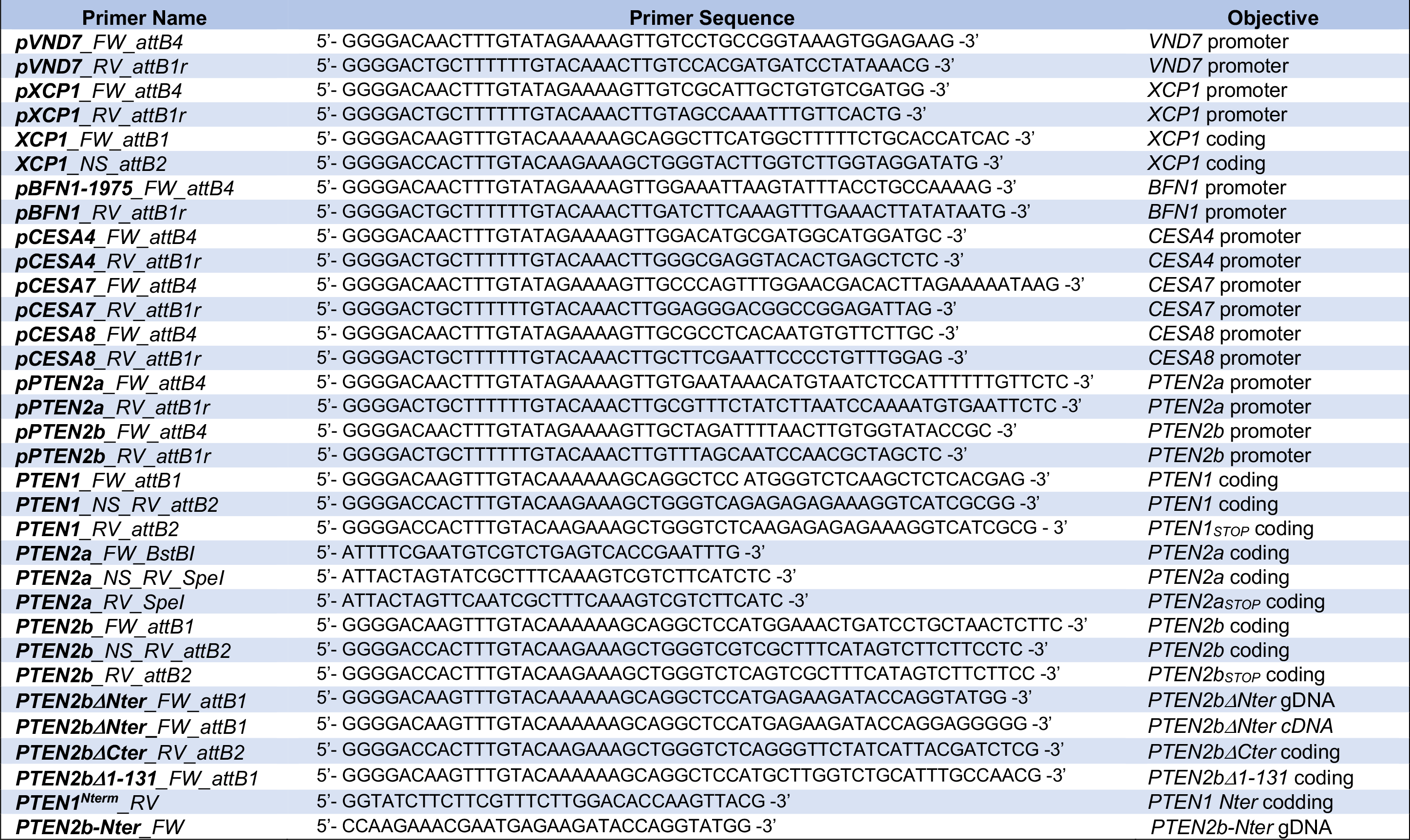
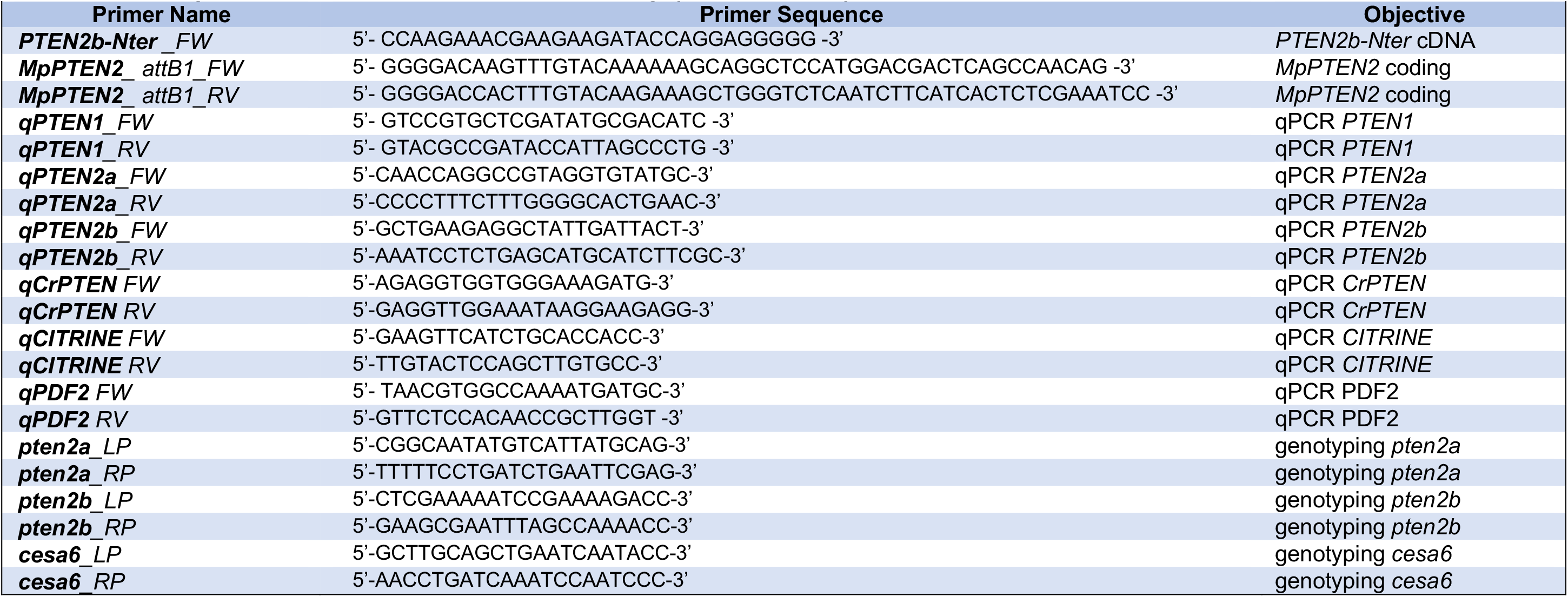
Primers used in this study.

**Supplementary Table 3:**
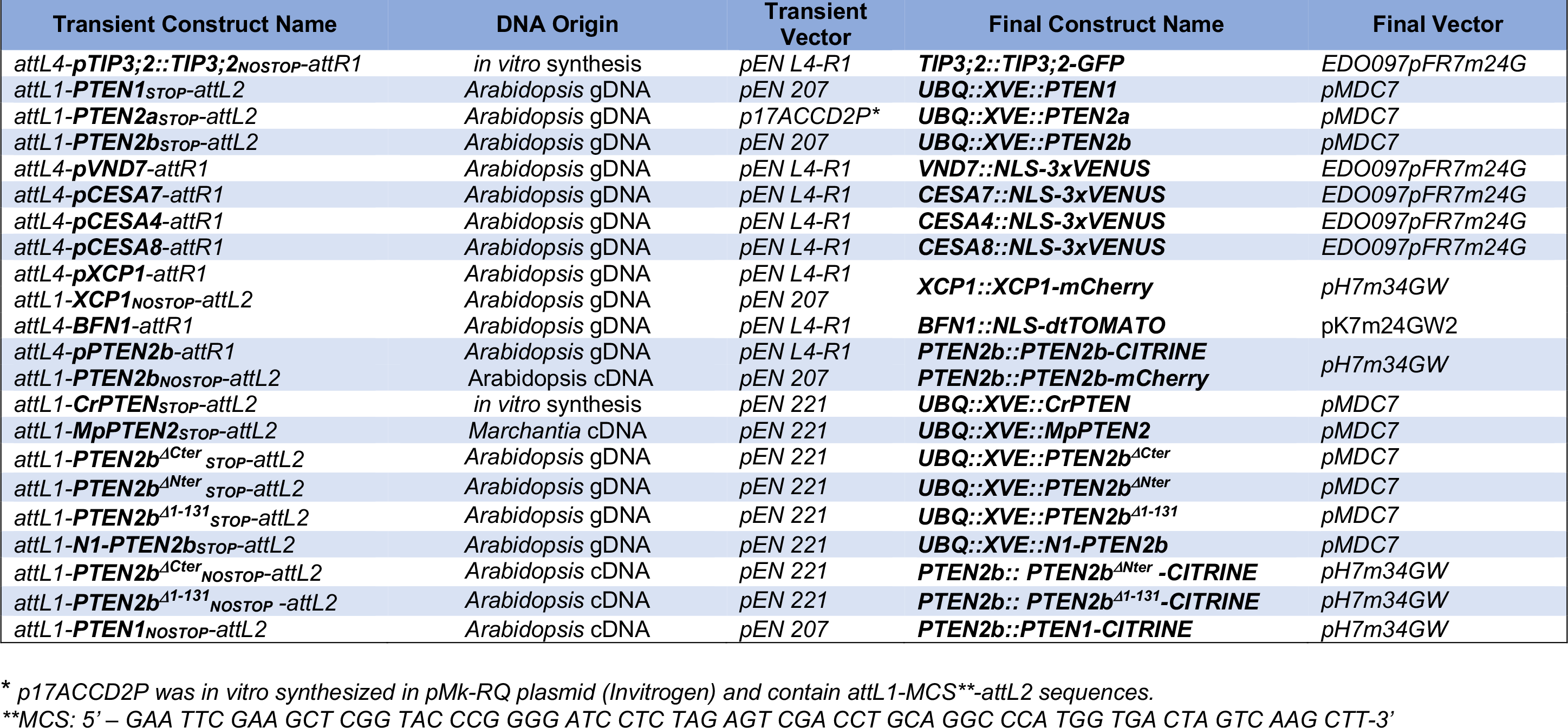
Constructs generated in this study.

| Transient Construct Name | DNA Origin | Transient Vector | Final Construct Name | Final Vector |
| --- | --- | --- | --- | --- |
| *attL4-pTIP3;2::TIP3;2<sub>NOSTOP</sub>-attR1* | *in vitro* synthesis | pEN L4-R1 | **TIP3;2::TIP3;2-GFP** | EDO097pFR7m24G |
| *attL1-PTEN1<sub>STOP</sub>-attL2* | *Arabidopsis* gDNA | pEN 207 | **UBQ::XVE::PTEN1** | pMDC7 |
| *attL1-PTEN2a<sub>STOP</sub>-attL2* | *Arabidopsis* gDNA | p17ACCD2P* | **UBQ::XVE::PTEN2a** | pMDC7 |
| *attL1-PTEN2b<sub>STOP</sub>-attL2* | *Arabidopsis* gDNA | pEN 207 | **UBQ::XVE::PTEN2b** | pMDC7 |
| *attL4-pVND7-attR1* | *Arabidopsis* gDNA | pEN L4-R1 | **VND7::NLS-3xVENUS** | EDO097pFR7m24G |
| *attL4-pCESA7-attR1* | *Arabidopsis* gDNA | pEN L4-R1 | **CESA7::NLS-3xVENUS** | EDO097pFR7m24G |
| *attL4-pCESA4-attR1* | *Arabidopsis* gDNA | pEN L4-R1 | **CESA4::NLS-3xVENUS** | EDO097pFR7m24G |
| *attL4-pCESA8-attR1* | *Arabidopsis* gDNA | pEN L4-R1 | **CESA8::NLS-3xVENUS** | EDO097pFR7m24G |
| *attL4-pXCP1-attR1* | *Arabidopsis* gDNA | pEN L4-R1 | **XCP1::XCP1-mCherry** | pH7m34GW |
| *attL1-XCP1<sub>NOSTOP</sub>-attL2* | *Arabidopsis* gDNA | pEN 207 |  |  |
| *attL4-BFN1-attR1* | *Arabidopsis* gDNA | pEN L4-R1 | **BFN1::NLS-dtTOMATO** | pK7m24GW2 |
| *attL4-pPTEN2b-attR1* | *Arabidopsis* gDNA | pEN L4-R1 | **PTEN2b::PTEN2b-CITRINE** | pH7m34GW |
| *attL1-PTEN2b<sub>NOSTOP</sub>-attL2* | *Arabidopsis* cDNA | pEN 207 | **PTEN2b::PTEN2b-mCherry** |  |
| *attL1-CrPTEN<sub>STOP</sub>-attL2* | *in vitro* synthesis | pEN 221 | **UBQ::XVE::CrPTEN** | pMDC7 |
| *attL1-MpPTEN2<sub>STOP</sub>-attL2* | *Marchantia* cDNA | pEN 221 | **UBQ::XVE::MpPTEN2** | pMDC7 |
| *attL1-PTEN2b<sup>ΔCter</sup><sub>STOP</sub>-attL2* | *Arabidopsis* gDNA | pEN 221 | **UBQ::XVE::PTEN2b<sup>ΔCter</sup>** | pMDC7 |
| *attL1-PTEN2b<sup>ΔNter</sup><sub>STOP</sub>-attL2* | *Arabidopsis* gDNA | pEN 221 | **UBQ::XVE::PTEN2b<sup>ΔNter</sup>** | pMDC7 |
| *attL1-PTEN2b<sup>Δ1-131</sup><sub>STOP</sub>-attL2* | *Arabidopsis* gDNA | pEN 221 | **UBQ::XVE::PTEN2b<sup>Δ1-131</sup>** | pMDC7 |
| *attL1-N1-PTEN2b<sub>STOP</sub>-attL2* | *Arabidopsis* gDNA | pEN 221 | **UBQ::XVE::N1-PTEN2b** | pMDC7 |
| *attL1-PTEN2b<sup>ΔCter</sup><sub>NOSTOP</sub>-attL2* | *Arabidopsis* cDNA | pEN 221 | **PTEN2b:: PTEN2b<sup>ΔNter</sup> -CITRINE** | pH7m34GW |
| *attL1-PTEN2b<sup>Δ1-131</sup><sub>NOSTOP</sub> -attL2* | *Arabidopsis* cDNA | pEN 221 | **PTEN2b:: PTEN2b<sup>Δ1-131</sup> -CITRINE** | pH7m34GW |
| *attL1-PTEN1<sub>NOSTOP</sub>-attL2* | *Arabidopsis* cDNA | pEN 207 | **PTEN2b::PTEN1-CITRINE** | pH7m34GW |

\* p17ACCD2P was *in vitro* synthesized in pMk-RQ plasmid (Invitrogen) and contain attL1-MCS\*\*-attL2 sequences.
\*\*MCS: 5' – GAA TTC GAA GCT CGG TAC CCG GGG ATC CTC TAG AGT CGA CCT GCA GGC CCA TGG TGA CTA GTC AAG CTT-3'

